# The division of amyloid fibrils – Systematic comparison of fibril fragmentation stability by linking theory with experiments

**DOI:** 10.1101/506386

**Authors:** David M. Beal, Magali Tournus, Ricardo Marchante, Tracey Purton, David P. Smith, Mick F. Tuite, Marie Doumic, Wei-Feng Xue

## Abstract

The division of amyloid protein fibrils is required for the propagation of the amyloid state, and is an important contributor to their stability, pathogenicity and normal function. Here, we combine kinetic nano-scale imaging experiments with analysis of a mathematical model to resolve and compare the division stability of amyloid fibrils. Our theoretical results show that the division of any type of filament is uniquely described by a set of three characteristic properties, resulting in convergence to self-similar length distributions distinct to each fibril type and conditions applied. By applying these results to profile the dynamical stability towards breakage for four different amyloid types, we reveal particular differences in the division properties of disease-related amyloid formed from alpha-synuclein compared with non-disease associated model amyloid, the former showing lowered intrinsic stability towards breakage and increased likelihood of shedding smaller particles. Our results enable the comparison of protein filaments’ intrinsic dynamic stabilities, which are key to unravelling their toxic and infectious potentials.

## INTRODUCTION

Amyloid fibrils, proteinaceous polymers with a cross-beta core structure, represent an important class of bio-nanomaterials ^1-2^. They are also important biological structures associated with devastating human diseases such as Alzheimer’s disease, Parkinson’s disease, Creutzfeldt-Jakob disease (CJD), systemic amyloidosis and type 2 diabetes ^3^, as well as vital biological functions such as adhesion and biofilm formation, epigenetic switches, and hormone storage (e.g. ^1-2, 4-9^). Division of amyloid fibrils, which can manifest *in vitro* in amyloid nano-materials or *in vivo* in disease associated or functional amyloid aggregates, is mediated by mechanical agitation, thermal stress, chemical perturbation or chaperone catalysis. It is a crucial step in the lifecycle of amyloid (**Fig. 1a**) ^10^, and enables the propagation of the amyloid protein conformation and biological information encoded therein. However, it is not understood why amyloid division processes give rise to varied biological impacts ranging from normal propagation of functional amyloid assemblies to large inert structures or the creation of molecular species involved in disease, e.g. small cytotoxic amyloid species and infective prions, which are transmissible amyloid particles.

**Figure 1.**
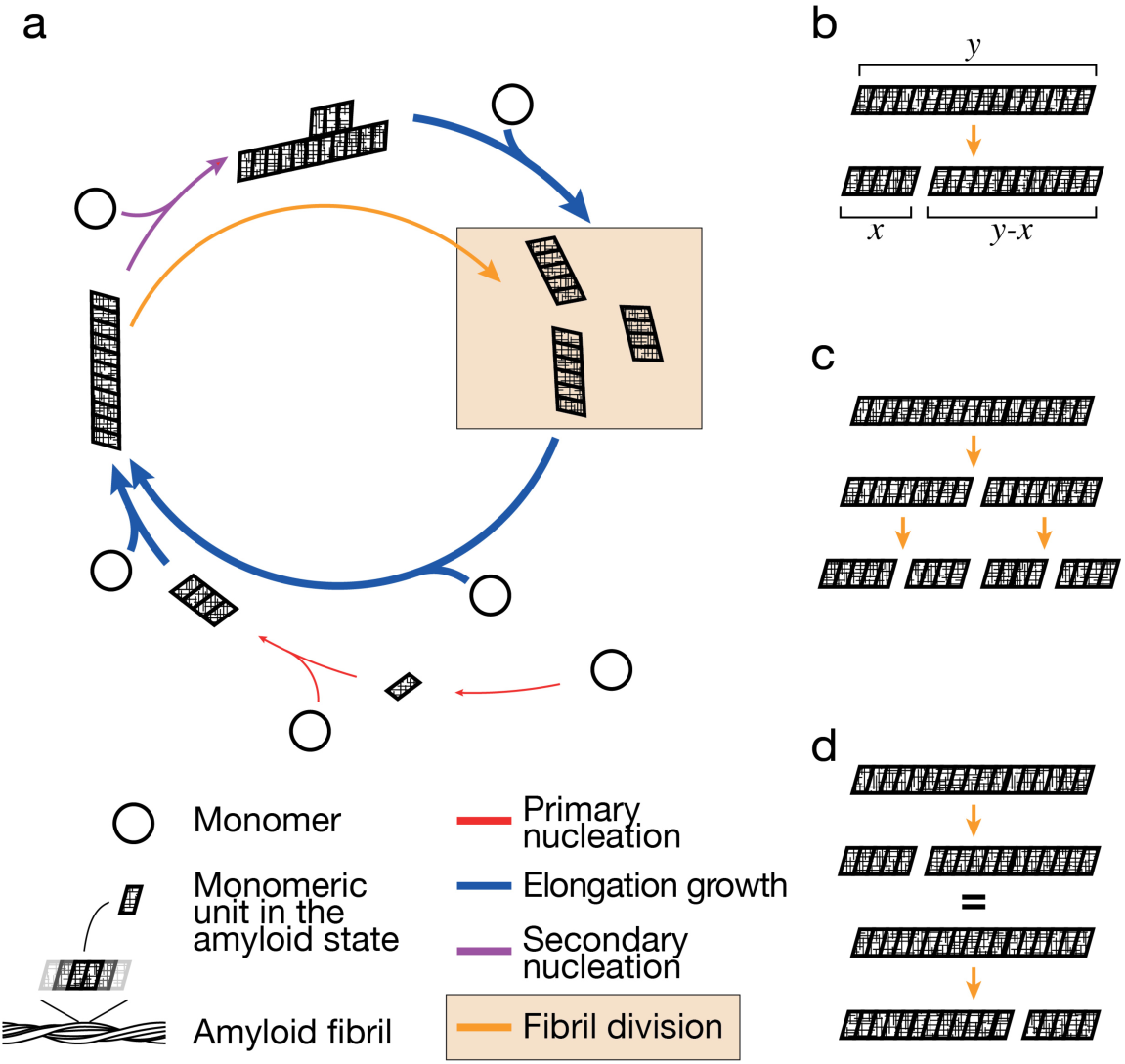
Schematic illustration of fibril division in the amyloid lifecycle. (a) The lifecycle of amyloid assembly where soluble monomeric protein (circles) are converted into the amyloid state with a cross-β conformation (the parallelograms). The coloured arrows represent the four main processes in amyloid assembly: primary nucleation (red), which may occur as homogeneous nucleation in solution or heterogeneous nucleation at interfaces; secondary nucleation (purple), which may occur as heterogeneous nucleation at surfaces presented by preformed aggregates; elongation growth at fibril ends (blue); and fibril division through for example fibril fragmentation or breakage (yellow, yellow box). The arrows may represent consecutive reversible steps and the thickness of the arrows symbolizes the relative rates involved in the processes. (b) A simple model of fibril division, where a given parent fibril particle of length y divide to give rise to two daughter fibril particles of size x and y-x. (c) The division model assumes that each mother fibril particle divides into exactly two daughter particles at each microscopic reaction step. (d) The division model assumes that the division rate for each microscopic step is identical as long as the resulting two particles have the same size.

The stability of these fibrillar bio-polymers are also important to the understanding of protein misfolding associated with disease progression and biological roles of functional amyloid assemblies (e.g. ^11^). In terms of disease association, there is much debate as to how amyloid aggregates are associated with cellular toxicity, with evidence of both prefibrillar oligomers and fibrillar species ^12-13^ giving rise to disease related phenotypes. While it is hypothesised that all proteins can undergo conversion into an amyloid state ^14^, why most proteins do not form amyloid under physiological conditions or produce amyloid particles that are non-toxic, non-transmissible or non-disease associated is not clear. In this debate, it has been suggested that fibrils are not merely the end product of amyloid aggregation, but rather elicit profound biological responses through fibril fragmentation and oligomer shedding ^12^, which may be the consequences of amyloid fibrils division due to lack of fibril stability.

Amyloid fibrils have remarkable physical properties, such as their tensile strength comparable to that of steel and an elasticity similar to spider silk ^15^. As proteinaceous polymers, they also offer the potential for modification by rational design, which makes them an ideal target for the development of biologically compatible nanomaterials ^2, 7, 16-17^. This interest in amyloid as a bio-nanomaterial has led to a search for proteins and peptides which can undergo conversion into a stable amyloid conformation while lacking the properties that associate them with toxicity, infectivity and disease. Though the precise properties that associate some amyloid to disease or to biological function are not resolved, the potential for different morphologies (sometimes referred to as ‘strains’) to elicit different results ^11, 18-19^ could be attributed to the stability of amyloid fibrils towards division or their mechanical properties ^20-21^. Thus, the stability of amyloid fibrils is an important factor which modulates their biological function of amyloid and potential as a nanomaterial.

The kinetics of the nucleated growth of amyloid fibrils are profoundly influenced by secondary processes such as fibril fragmentation/breakage ^22-23^ and secondary surface nucleation ^24-25^ (**Fig. 1a**). These processes determine the rate of the exponential growth phase of amyloid assembly alongside with elongation growth at fibril ends ^22, 26^. As one of the key secondary processes, fibril fragmentation stands out compared to the other three main processes (**Fig. 1a**) in that it reduces aggregate size at the same time as it increases the number of aggregates ^21^. In this aspect, fibril fragmentation results in the division of amyloid fibrils analogous to a microbial or cellular division processes. Resistance towards fibril division by fragmentation is linked to the mechanical stability of amyloid fibrils, which has implications for both the development of nanomaterials and on the understanding of amyloid disease associated biological processes.

The division of amyloid polymers into small more infective particles, either through environmental perturbations or through catalysis by molecular chaperones, is key to the spreading of prion phenotypes ^20, 27^. In addition, the smaller particles generated by fibril fragmentation show enhanced cytotoxicity when compared with the larger parent fibrils ^21^, likely due to an greater propensity to interact with cell membranes, entering cells by endocytosis, interacting with the lysosome and inducing cytotoxicity by disrupting proteostasis ^20, 28-31^. The stability of amyloid fibrils towards division is, therefore, an important characteristic of amyloid fibrils that must be considered if we are to understand the biological activity and nanomaterial properties of amyloid. Because protein filaments formed from different precursors show a variety of suprastructures and size distributions (e.g. ^15, 18, 21, 32^), no unifying theory has been developed for the division of amyloid fibrils. As consequence, the stability towards division for different types of amyloid fibrils with varied suprastructures that ranges from inert network of long filaments to infectious particles is yet to be systematically measured, determined and compared.

The mechanism and the rate of division for amyloid filaments has been subjected to theoretical considerations ^22-23, 33-34^ and experimental investigations involving fibril fragmentation promoted by mechanical perturbations ^22, 35-36^. The fragmentation of protein filaments is a length dependent process whereby longer particles break more easily than short ones. This length-dependent breakage of amyloid fibrils can follow a strong, non-linear dependence where longer fibrils are progressively less stable towards breakage per monomeric unit relative to their shorter counterparts ^35^. Thus, the fibrils’ resistance to division, and in turn the inherent stability of the fibrils, is an important and measurable property ^35^ that will help to rationalise phenomena such as prion strains, polymorphism, transmission, amyloid toxicity, biofilm formation and epigenetic regulation (e.g. ^11, 20-21, 27, 37-43^) and leading to a better understanding of amyloid-associated diseases.

We have previously shown that the time evolution of amyloid fibril length distributions obtained by nano-scale atomic force microscopy (AFM) imaging contain valuable information on the rate, length-dependence and position-dependence of fibril fragmentation that can be extracted ^35^. However, since fibril division is itself a strongly length-dependent process, systematic comparison of the stability of fibrils towards division and their division rates has been hampered by the varied length distributions of different types of amyloid fibrils. Currently, the links between data and theory that would allow direct comparison of the fibrils’ division propensities are also missing. Here, we have developed an analytical approach that enables direct determination of the dynamic stability of amyloid fibrils towards division from fibril length distributions. We have developed a new theory on amyloid fibril division that shows how the division mechanism of amyloid fibrils, and their stability towards division dictates the exact shape of the resulting length distributions. We then established an analytical method to extract a set of unique and intrinsic properties of the fibril division processes from image data of fibrils undergoing division experimentally promoted by mechanical perturbation. Demonstrating the utility of our combined experimental and theoretical approach, we determined and compared the division of fibril samples formed from human α-synuclein (α-Syn) associated with Parkinson’s disease with fibrils formed from β-lactoglobulin (β-Lac) and lysozyme (Lyz). We have also reanalysed and compared previously published fibril fragmentation data of β_2_-microglobulin (β_2_m) under the same mechanical perturbation regime^35^. Comparison of the dynamic stability of these fibrils types of different origin revealed different division properties, with fibrils formed from the human Parkinson’s disease-associated α-synuclein being the least overall stable and prone to generate small sub 100 nm particles that may possess enhanced cytotoxic and prion-like infectious potential ^44^. The ability to assess and compare the division properties of amyloid fibrils, enumerated as parameters extractable from experimental data, enables the prediction of an amyloid’s propensity to generate toxic and infectious particles, and therefore has a significant impact on the understanding of their roles in biology, in diseases, and their application as a functional bio-nanomaterial.

## RESULTS

### Amyloid fibrils of diverse suprastructures and length distributions fragment to different extents upon mechanical perturbation

To demonstrate that the fibril division rates, indicative of their dynamic stability towards division, can be assessed and compared for amyloid fibrils with diverse suprastructures and length distributions, we first collected experimental AFM image data sets of amyloid fibrils, pre-formed from different precursors, undergoing division through fragmentation promoted by mechanical stirring. These experiments were designed to isolate the fibril division processes from other growth processes and to generate data that contain sufficient quality and quantity of information on the division of fibril particles under identical mechanical perturbation regimes to enable comparison. Here, we chose to investigate the human disease-associated amyloid system α-Syn alongside bovine β-Lac and chicken egg white Lyz as biophysical model systems not directly related to human disease. Samples were formed containing long-straight fibrils from these three proteins *in vitro*. Lyz and β-Lac were both converted to their fibrillar amyloid form by heating under acidic conditions (pH 2.0), commonly used conditions for the assembly of these proteins *in vitro*. α-Syn fibrils were prepared from freshly purified recombinant α-Syn monomers at 37°C under physiological pH. For each fibril sample, 500 *µ*l of 120 *µ*M monomer equivalent fibril solutions in their respective fibril forming buffer were then stirred at 1000 rpm by a 3 x 8mm magnetic stirrer bar in a 1.5ml glass chromatography vial using the same mechanical perturbation method as previously reported ^35^ using an Ika Squid stirrer plate with a digital display. The *in vitro*-formed fibril samples were initially dispersed by 5-10 min of stirring and were subsequently deposited onto freshly cleaved mica surfaces and imaged by AFM (**Fig. 2** left most column).

**Figure 2.**
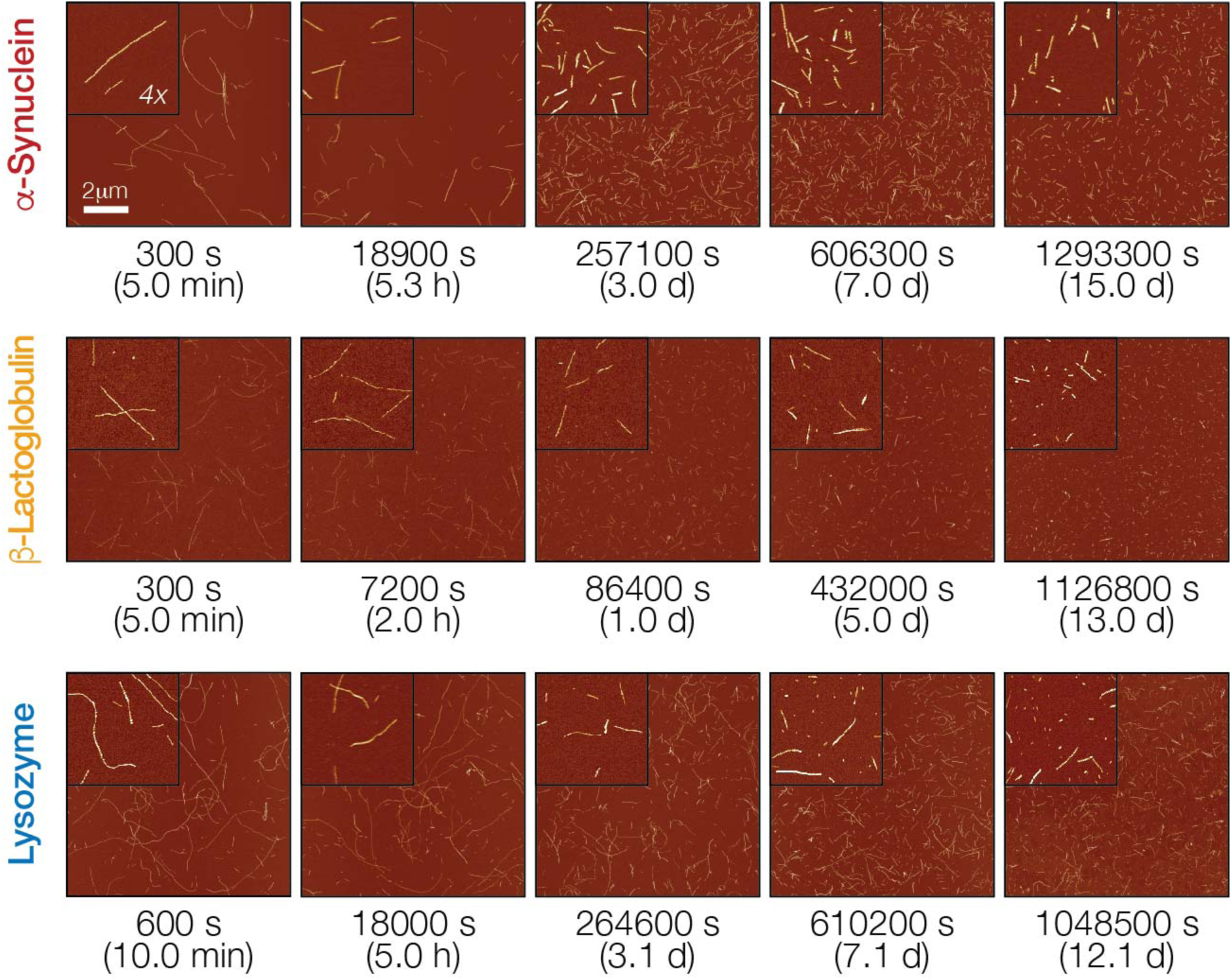
AFM imaging of amyloid fibrils undergoing division through fragmentation promoted by mechanical stirring. Hen egg white Lyz, bovine milk β-Lac, and human α-Syn amyloid fibril samples (all 120 *µ*M monomer equivalent concentration) were stirred for up to 15 days. Samples were taken out periodically, deposited on mica and imaged using AFM. Typical AFM images representing 10×10 μm surface areas are show together with 4x magnified insets. The scale bar represents 2 μm in all images.

As seen in the leftmost column of images in **Fig 2.**, the initial samples after brief stirring to disperse the fibril particles show long, straight, elongated, unbranched nano-structures expected for amyloid fibrils. However, whereas Lyz and α-synuclein form fibrils that exhibit more flexibility and curvature, β-Lac forms comparably straighter, more rigid assemblies consistent with previous observations (e.g. ^15, 36, 45-46^). Importantly, all of the samples showed well-dispersed fibril particles that can be individually measured after the brief stirring treatment, as the samples did not show strong propensity for clumping on the surface substrates.

The samples were then continuously stirred for up to 15 days and 1-5 *µ*l samples (see materials and methods) were taken out periodically and imaged using AFM to visualise their fragmentation under mechanical perturbation (**Fig. 2**). For each sampling time-point, an identical AFM specimen preparation procedure was used for each amyloid type, and 20 *µ*m x 20 *µ*m surface areas were imaged at 2048 x 2048 pixels resolution in order to enable quantitative analysis of individual fibril particles as previously described ^47^. In total, fragmentation of two independent fibril samples was followed for each fibril type, and 171 images with at least 300 particles for each sample and time point were analysed, giving a total dataset containing physical measurements of more than 220,000 individual fibril particles for the three amyloid types (**Supplementary Table S1**).

Quantitative single-particle measurements of fibril length and height distributions (**Fig. 3**, leftmost column corresponding to images in **Fig. 2**. leftmost column) reveal that the fibrils have substantially different initial dimensions. Analysis of their height distributions show that the initial fibril heights, indicative of the width of the fibrils, are around 7 nm for α-Syn fibrils, and around 3 nm for both β-Lac and Lyz fibrils. The initial length distributions for the different fibril types were also dissimilar, with both Lyz and α-Syn forming fibrils of up to ∼10 *µ*m in length whereas β-Lac formed shorter particles with lengths of up to ∼2 *µ*m under the conditions employed.

**Figure 3.**
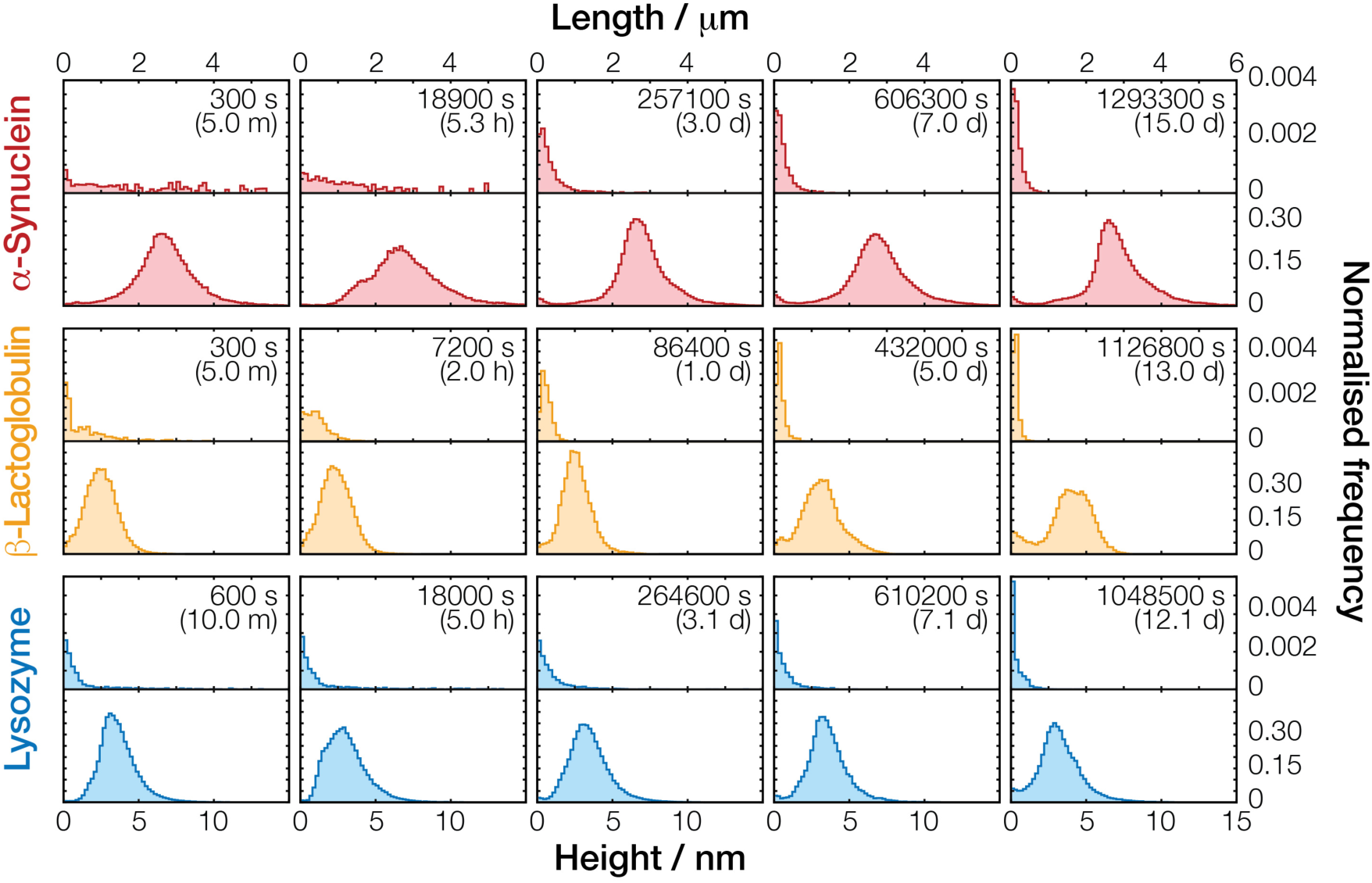
Fibril length and height distributions extracted from AFM images of the fibril undergoing fragmentation by mechanical perturbation. Normalised length (upper row of each sample) and height (lower row of each sample) distributions of fibril particles corresponding to the same AFM images in **Fig. 2** are shown as histograms. The histograms are shown using the same length and height scales, respectively, for comparison.

Qualitative inspection of the AFM images over the duration of the experiment (**Fig. 3**) showed that the amyloid fibrils were fragmented into much smaller particles under the applied mechanical perturbation (**Fig. 2** and **Fig. 3**) as expected. However, the rate of division and shortening of the particles’ lengths was seen to differ between the three different fibril types analysed (**Fig. 3** and **Supplementary Fig. S1**). Analysis of the time evolution of the fibril height and length distributions obtained by quantification of individual particles in the AFM images over the course of the experiment confirmed that fibril fragmentation did not cause detectable changes in fibril morphology and fibril width through lateral association and dissociation. Average fibril heights in the AFM images, indicative of fibril widths, remained consistent throughout the experiment for Lyz and α-Syn, the same were also largely observed for β-Lac, with the exception that a small second population of taller polymers at the very end of the fragmentation time-course after 432000 s were exhibited (height graphs in **Fig. 3** and **Supplementary Fig. S1**). Hence division of the fibrils under mechanical perturbation applied has resulted in a shortening of average fibril length.

To confirm that the changes in fibril length by fibril division did not cause disaggregation or release of monomer/small oligomers (e.g. dimers), we next determined the residual monomer concentration of the samples. For each fibril type, aggregates were pelleted by centrifugation (75k rpm, 15 min) after fragmentation time-course and the presence of monomer in the supernatants was quantified by SDS-PAGE. The comparison between the initial samples and those fragmented over two weeks showed no large changes in the protein composition of the supernatants, with differences of less than 2% for all amyloid systems analysed (Lyz: 1.4%, β-Lac: <1%, and α-Syn: 1.3%, **Supplementary Fig. S2**). These data confirmed that the time-dependent imaging experiments we carried out pertain almost exclusively to the fibril division processes along the length of the pre-formed fibrils, and therefore, contain valuable information on their division rates and their stability towards division.

### Time evolution of fibril length distributions converges to time-independent, characteristic, self-similar length distribution shapes

The fibril samples formed from different protein precursors have different initial length distributions (as seen in **Fig. 2** and **Fig. 3**). However, fibril division is itself a strongly length-dependent process ^35^ as short fibril particles will be more resistant towards division compared to longer particles, irrespectively of any differences in the intrinsic stability of the different fibril types towards division. Therefore, to compare the stability of amyloid fibrils with different suprastructures and length distributions towards division, a new approach to extract information intrinsic to each fibril type independent of their experimentally different initial length distributions must be developed. Consequently, in parallel with the experiments described above, we mathematically analysed the division equation of amyloid fibrils so that key information on the stability of amyloid fibrils towards division could be resolved. We first describe mathematically the division of amyloid fibrils using a continuous framework based on the partial differential equation (PDE) Eq. (1). Since the number of monomers inside a fibril observed in the image data is large, typically in the order of 10^2^ or more, we assumed continuous variables *x* and *y* that correspond to the length of fibrils (for example as defined in **Fig 1b** where y is the length of the parent fibril and x is the length of one of the daughter fibrils). This approach has the advantage that the infinite set of ordinary differential equations (ODEs) normally used to describe the length-dependent division processes (e.g. ^22-23, 35^) can now be collapsed into a single continuous PDE that can be treated analytically (see Supplemental Information for details). Denoting *u(t,x)* as the distribution of fibrils of length *x* at time *t* in number concentration units (e.g. Molar units), Eq. (1) is the mathematical translation of the pure division model described by the schematics in **Fig.1b-d**, where we assume any parent fibril can divide into two daughters, and the end-end reattachment rate of daughter fibrils is negligibly slow ^33^:

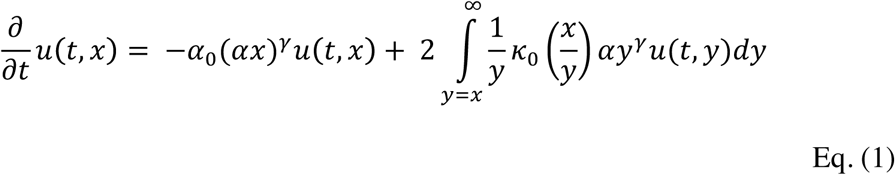

In Eq. (1), 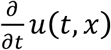 denotes the time (*t*) evolution of the concentration of fibrils with length *x*. Here, we model the total division rate constant of fibrils of size *x* using the power law *α*_0_(*αx*)^*γ*^, which we denote as *B*(*x*) ^33^ (see Supplementary Information), where *α*_0_ is a constant unit reference we set to 1 s^-1^. The first term in Eq (1), therefore, denotes the rate of loss of fibrils with length *x* by division into smaller fibrils. The probability that after dividing, a given parent fibril of length *y* gives rise to a daughter fibril fragments of length *x* and *y-x* depends on the ratio of the lengths (*x/y*) ^35^ and is given by the probability density function 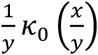. The second integral term in Eq. (1), therefore, denotes the total gain of fibrils with length *x* by division of all fibrils with length *y* that are larger than *x*. Interestingly, Eq. (1) describes a fundamental division process that is mathematically analogous to the division of molecules, macroscopic materials and cells ^48-49^, and we have mathematically proven that its behaviour is entirely and uniquely dictated by three properties: *α* that describes the magnitude of the division rate constant, *γ* that describes the fibril length dependence of the division rate constant, and *κ*_*0*_ that describes the probability of division at any given position along a fibril, also called the fragmentation kernel ^50^. We then proceeded to solve Eq. (1) analytically with regard to *α, γ* and *κ* using theoretical results shown in ^49^ and ^50^ (see Supplementary information). From our solution, we note four key predictive insights that emerged from our analysis (**Fig. 4**).

**Figure 4.**
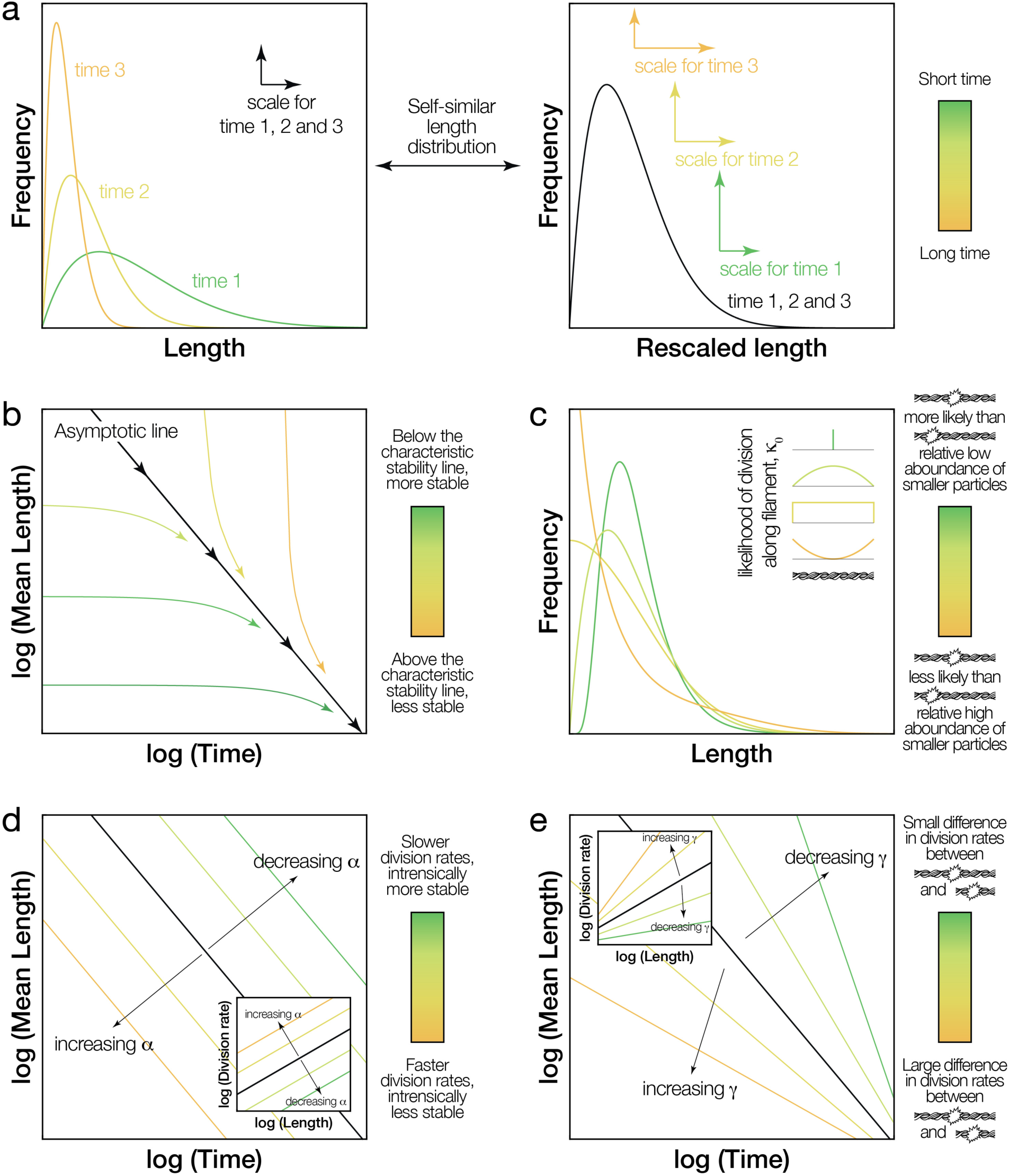
Illustration of the key insights emerging from the mathematical analysis of the division model. The behaviour of the division equation Eq. (1) is entirely and uniquely dictated by a set of three properties: α, γ, and κ_0_. Several key predictive insights emerged from the analytical solution of Eq. (1) with regard to these three properties. (a) The three example length distributions in the left panel can be rescaled to show the same distribution shape in the right panel, illustrating the concept of self-similar length distributions. (b) After a sufficiently long time where the self-similar length distribution shape is reached, the reduction in the average length of the fibril length distribution can be described as a power law versus time. The decay of mean length of a sample is predicted to tend towards a straight line, the asymptotic line, when plotted on a log-log plot with the slope of the line representing −1/γ (black line in b, d and e). The stability line with mean fibril lengths also does not depend on the initial length distribution (coloured lines in b). (c) The self-similar length distribution shape contains information about κ_0_, which describes how likely a fibril will divide in the middle versus shedding a small fragment from the edge. A κ_0_ indicative of fibril types that are more likely to divide in the middle will result in fibril length distributions with a distinct peak and low relative population of small fragments (dark green and light green curves). In contrast, κ_0_ indicative of fibril types and conditions that promote equal likelihood of division along the fibril or even favour shedding of small fragments from fibril edges will result in self-similar fibril length distributions that have a larger relative population of small fibril fragments (yellow and orange curves) compared to κ_0_ values favouring division in the centre of the fibrils. (d) and (e) illustrates how the black asymptotic line describing the decay of fibril lengths in (a) is dictated by the parameters α and γ, respectively. For each panel, the colour bar to the right illustrate the different properties associated with the colours in the panel (e.g. division in the centre vs. at the edge of a fibril for panel c, and division of a long vs. a short fibril in panel e)

Firstly, we note that given enough time, the decay of the average fibril length will converge to the same rate independently of the initial fibril length distributions. This result comes from that after a sufficiently long time, the reduction of average length of the fibril length distribution can be described as a power law versus time (Eq. 2, see Supplementary Information):

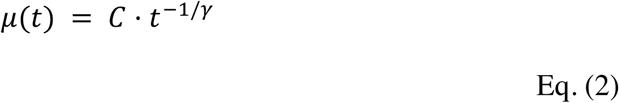

where *C* is a constant. As seen in Eq. (2), the experimentally observable average length of a sample, *μ*(*t*), is predicted to tend towards a straight line when plotted on a log-log plot with the slope of the line representing −1/*γ* (Eq. 2, black line in **Fig. 4b**) because the long-time behaviour of Eq. (1) can be described as an elegant power law.

Secondly, we note that given enough time, the fibril length distribution will converge to the same shape independently of the initial state of the fibril length distribution. After a sufficiently long time (*t* ≫ *t*_0_), the distribution of fibril-lengths tends towards a time-independent distribution shape, *g*(*x*_*g*_), that scales only with *t* and *γ*, but does not depend on the initial length distribution (Eq. (3) and Supplementary Information).

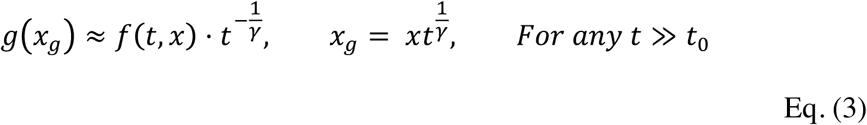

where *f(t, x)* are experimentally measured length distributions. This point is of key importance for characterising and predicting fibril division processes because it establishes that for any fibril type under certain conditions: 1) a distinct fibril length distribution shape (**Fig. 4a**) will be reached independently of the initial fibril length distribution, and 2) the length distribution and the average length will shrink as function of time in a predictive manner as fibrils continue to divide (e.g. the black line in **Fig. 4b** for the mean length) but the shape of the distribution will not change as function of time, i.e. the length distribution can be rescaled to the same *g*(*x*_*g*_) using Eq. (3) at any time *t* along the black line in **Fig. 4b** if *t* is sufficiently large. We refer to the distributions with the scaling property and shape invariance property as ‘self-similar length distributions’ (**Fig. 4a**).

The existence of a self-similar length distribution that is initial length distribution-independent and shape invariant over time, as well as the predictable decay of fibril lengths as fibrils divide (e.g. the reduction of the average length in **Fig. 4b**) can be seen as a characteristic behaviour specific to individual fibril types under distinct conditions. This fibril division behaviour can, therefore, be classed as a type of intrinsic dynamic stability of the fibrils. One way to visualise this property is shown in **Fig. 4b** represented by the black line, here referred to as the fibril type’s ‘asymptotic line’ under the conditions applied. Any fibril populations above this line are relatively unstable and will rapidly divide, pushing the average length towards the line (yellow coloured near-vertical arrows showing rapid decay of unstable fibril lengths), while any fibril populations below this line are comparatively stable or metastable and will only slowly evolve towards the line through division (green coloured near-horizontal arrows showing slow decay of stable fibril lengths towards the black line). Importantly, this result also indicates that the dynamic stability of fibrils towards division represented by the asymptotic line: 1) can be determined from experimental data, 2) is intrinsic to fibril type and conditions applied, and 3) can be compared independently of varied starting fibril length distributions, if the characteristic self-similar length distributions that contain information about the intrinsic dynamic stability of the fibrils is reached (e.g. the asymptotic line is reached in an experiment running for sufficiently long length of time).

Thirdly, we note that the probability of division in the centre of a fibril as compared to shedding of small particles from fibril edge can be evaluated from the experiments. The self-similar length distributions contain information about κ_0_, which describes how likely a fibril will divide in the middle compared to shedding a small fragment from the edge. **Fig. 4c** shows how different self-similar fibril length distributions are indicative of different κ_0_ probability functions. As seen in **Fig. 4c**, κ_0_ indicative of fibril types that are more likely to divide in the middle will result in fibril length distributions with a distinct peak and low relative population of small fragments. In contrast, κ_0_ indicative of fibril types and conditions that promote equal likelihood of division along the fibril or even favouring the shedding of fragments from fibril edges will result in self-similar fibril length distributions that have large relative population of small fibril fragments that may possess enhanced cytotoxic and/or infective potential compared to κ_0_ favouring division in the centre of the fibrils.

Finally, the dynamic stability of fibrils towards division, their propensity to break at different lengths, can be determined. The first order division rate constant *B*(*x*) = *α*_0_(*αx*)^*γ*^ that describes the division of the fibrils as a function of their length *x* can be directly evaluated from the self-similar length distribution shape and *γ* (see Eq. 2) when *t*≫*t*_*0*_ (see Supplementary information and Eq SI.21). Thus, the division rate constant *B(x)* can be determined from experimentally observing how fibril length distributions change with time when the self-similar fibril length distribution is obtained, and they are important parameters for defining and comparing the fibrils intrinsic dynamic stability towards division. The effect of different values of *α* and *γ* on fibril stability is visualised in **Fig. 4d** and **Fig. 4e** as characteristic the asymptotic line plotted in log-log plots of average length versus time. The enumeration of the asymptotic line described by *B(x)* will subsequently enable direct quantitative comparison of the fibrils’ stabilities towards division.

### The division properties of amyloid fibrils can be obtained from image data and their complex stability towards division can be compared

Applying the results of the mathematical analysis to the experimental AFM image data sets, the parameters *γ, α*, and the characteristic self-similar length-distributions *g*(*x*_*g*_) indicative of *κ*_*0*_ can be extracted and meaningfully compared as a measure of the fibrils’ intrinsic stability towards division. We first determined the *γ* values for each of the fibril types, by globally fitting a variant of Eq. (2) to the time evolution of average fibril length (see Materials and methods, **Fig. 5**). We also reanalysed previously published data set on β_2_m fibril fragmentation under the same mechanical perturbation conditions ^35^ using our new theoretical results above and included the reanalysis in the comparison.

**Figure 5.**
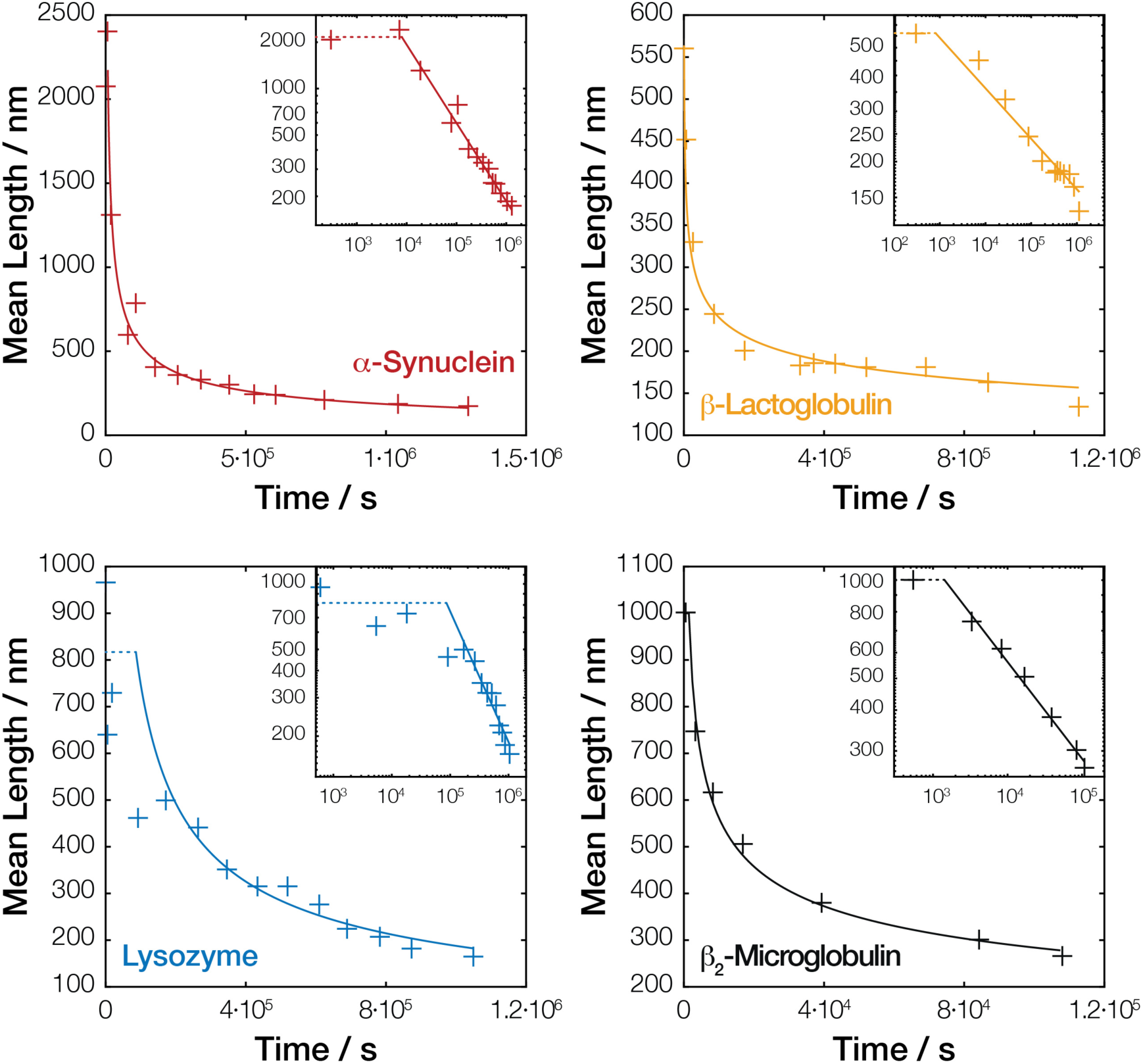
Fitting the fibril division model to fibril length decay data extracted from AFM images. The analytical solution of our division model shows the decay of average length as function of the gamma parameter in equation Eq. (2) and Eq. (4). Equation Eq. (4) was fitted to the decay of average fibril length during division for each of the fibril types analysed (including previously published data for β_2_m fragmentation under the same mechanical perturbation conditions ^35^). The solid fitted lines represent the time regime where the length distributions closely approached the stability line and the self-similar distribution shape where Eq. (2) is valid (Materials and Methods).

The constant *γ* relating to the fibril length dependence of the division rate constant was determined from least-squares fitting of our analytical result to the data (Materials and Methods). The elegant power law relationship (Eq. 2) parameterised with *γ* determined by global analysis were visualised on a log-log plot of mean fibril length vs. time in **Fig. 5**, together with the measured mean fibril lengths. The resulting *γ* values are listed in **Table 1**. A *γ* value of 1 would suggest that the division rate of fibrils is only dependent on the number of division sites per fibril, which is linearly related to the number of monomers in the fibrils and in turn to the length of the fibrils. However, the *γ* values for α-Syn, β-Lac and β_2_m are all significantly larger than 1, indicating highly length dependent microscopic division rates for division sites in these fibril types. Out of the four fibril types analysed, only the division of Lyz fibrils yielded a *γ* value closest to 1 suggesting the division rates for Lyz fibrils may only depend on the number of available division sites along the fibrils. As seen in **Fig. 5**, the later time points for all of our fibril types follow a straight line on the log-log plots (solid section of the fitted lines in **Fig. 5**), indicating that the self-similar length distributions, and hence the asymptotic line, were sufficiently reached in all cases. The analysis also revealed that all of the fibril types analysed approached the self-similar length distribution shapes in less than 5 hr, with the exception of the Lyz samples that reached the self-similar distribution in approximately 24 hr.

**Table 1.** Parameters from the division analysis of the different fibril types

| *Sample* | $\gamma \pm SE$ | $\alpha / nm^{-1} (\log \alpha \pm SE)$ | $B (100 nm) / s^{-1} (\log B \pm SE)$ | *Width / nm* |
| --- | --- | --- | --- | --- |
| *α-Syn* | $2.0 \pm 0.3$ | $2.6 \cdot 10^{-6} (-5.6 \pm 0.2)$ | $9.2 \cdot 10^{-8} (-7.0 \pm 0.3)$ | $6.8 \pm 0.6$ |
| *β-Lac* | $5.7 \pm 0.8$ | $1.8 \cdot 10^{-4} (-3.7 \pm 0.2)$ | $1.2 \cdot 10^{-10} (-9.9 \pm 0.8)$ | $3.0 \pm 0.5$ |
| *Lyz* | $1.7 \pm 1.0$ | $9.4 \cdot 10^{-7} (-6.0 \pm 0.9)$ | $2.0 \cdot 10^{-7} (-6.7 \pm 1.0)$ | $3.1 \pm 0.4$ |
| *$\beta_2m^*$* | $3.4 \pm 0.4$ | $5.6 \cdot 10^{-5} (-4.3 \pm 0.3)$ | $2.5 \cdot 10^{-8} (-7.6 \pm 0.4)$ | $5.4 \pm 0.6$ |

\* Reanalysis of data from Xue and Radford 2013<sup>35</sup>.

The *α* values were subsequently calculated (listed in **Table 1**) with equations Eq. (SI.21) using all of the fibril length distributions at time points post reaching the near-characteristic self-similar distribution shapes (represented by the solid lines in **Fig. 5**). Once both *α* and *γ* values have been extracted from the length-distribution data, the division rate constant *B(x)* can be obtained for fibrils of any length *x*. **Table 1** shows the division rate constant calculated for example fibrils of 100 nm. The asymptotic line for the fibrils types characterised by the division rate constant *B(x)* (**Fig. 6b**) or by fibril mean length (**Fig. 6a**) as function of time was also visualised and compared independently of initial fibril length, showing that α-Syn and Lyz fibrils fragments fastest at long times under the mechanical perturbation applied, suggesting that these fibrils were less stable then the β-Lac and β_2_m fibrils.

**Figure 6.**
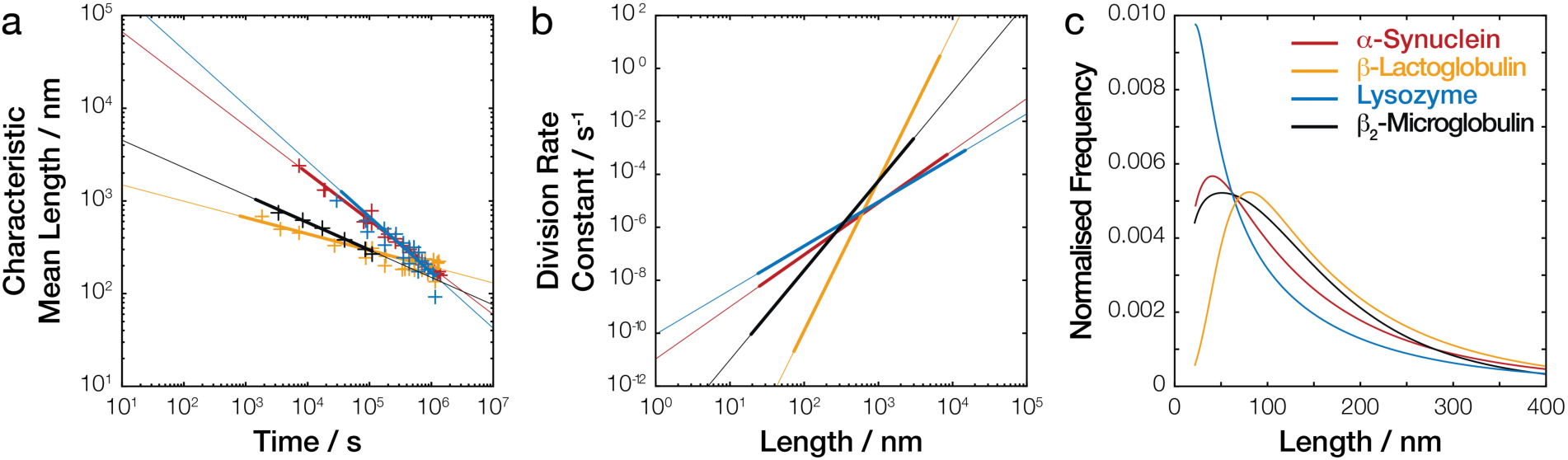
Comparing the stability towards division of different amyloid fibril types. The decay of mean lengths (a), the division rate constants as function of fibril length (b), and the self-similar length distribution shapes (c) for hen egg Lyz (blue), bovine milk β-Lac (yellow), human α-Syn (red) and human β_2_ m (black, data from Xue and Radford 2013 ^35^) amyloid fibril samples undergoing division by fibril fragmentation under mechanical perturbation. All curves were calculated using α, γ, and g(x_g_) obtained from our analysis of the experimental AFM images. In (a), the thicker portion of the lines denote the time range where the characteristic self-similar length distribution shape is observed in the imaging experiments (i.e. corresponding to the time regime represented by the solid fitted lines in **Fig. 5**), and crosses are the experimental data points that have closely reached the self-similar distribution shapes shown in the same plot. In (b), the thicker portion of the lines denote the range of fibril lengths observed experimentally on the AFM images. In (c), the distributions were calculated using self-similar distributions g(x_g_) in **Supplementary Fig. S3** after two weeks.

Next, we determined the shape of the self-similar length distributions for each fibril type by rescaling the experimental length distributions to *g*(*x*_*g*_) with Eq. (3) using the *γ* values obtained above. As with the evaluation of *α* values, only time points where the length distributions closely approached the self-similar length distribution (time points in the section represented by the solid lines in **Fig. 5**) where averaged to obtain *g*(*x*_*g*_) for each fibril type (**Supplementary Fig. S3**). **Fig. 6c** shows how the self-similar length distribution shapes compare with each other at extended times (2 weeks) when calculated using *g*(*x*_*g*_) (**Supplementary Fig. S3**) with Eq. (3). As seen in Fig 6c, Lyz fibrils have a tendency to produce high relative populations of small particles less than 100 nm long followed by α-Syn and then β_2_m. On the other hand, the division of β-Lac fibrils resulted in a lower relative population of small particles over the same long time scale used for the other fibril types.

Finally, to validate our model and the predictive power of our approach, we performed direct simulations of the fibril division time-course (**Fig. 7**) using only the individual sets of division parameters obtained for each of our fibril types. For each simulation, we used the initial experimental length distributions (dashed lines in **Fig. 7**) directly as the starting points for the simulations and solved the large set of ordinary differential equations describing the chemical master equation for the system ^35^ to see whether our analytical model was able to predict their full division behaviour and the time evolution of the fibril length distributions for each fibril type. As seen in **Fig. 7**, the result of the numerical simulations based on our results show remarkable agreement with the experimental data. This unequivocal result validated the fact that the set of three properties *γ, α*, and *κ*_*0*_ are indeed capable of fully and uniquely describing the complex amyloid division processes, and the enumeration of these properties yield valuable insights. Such insights allow meaningful comparison of the amyloid fibrils’ intrinsic stability towards division.

**Figure 7.**
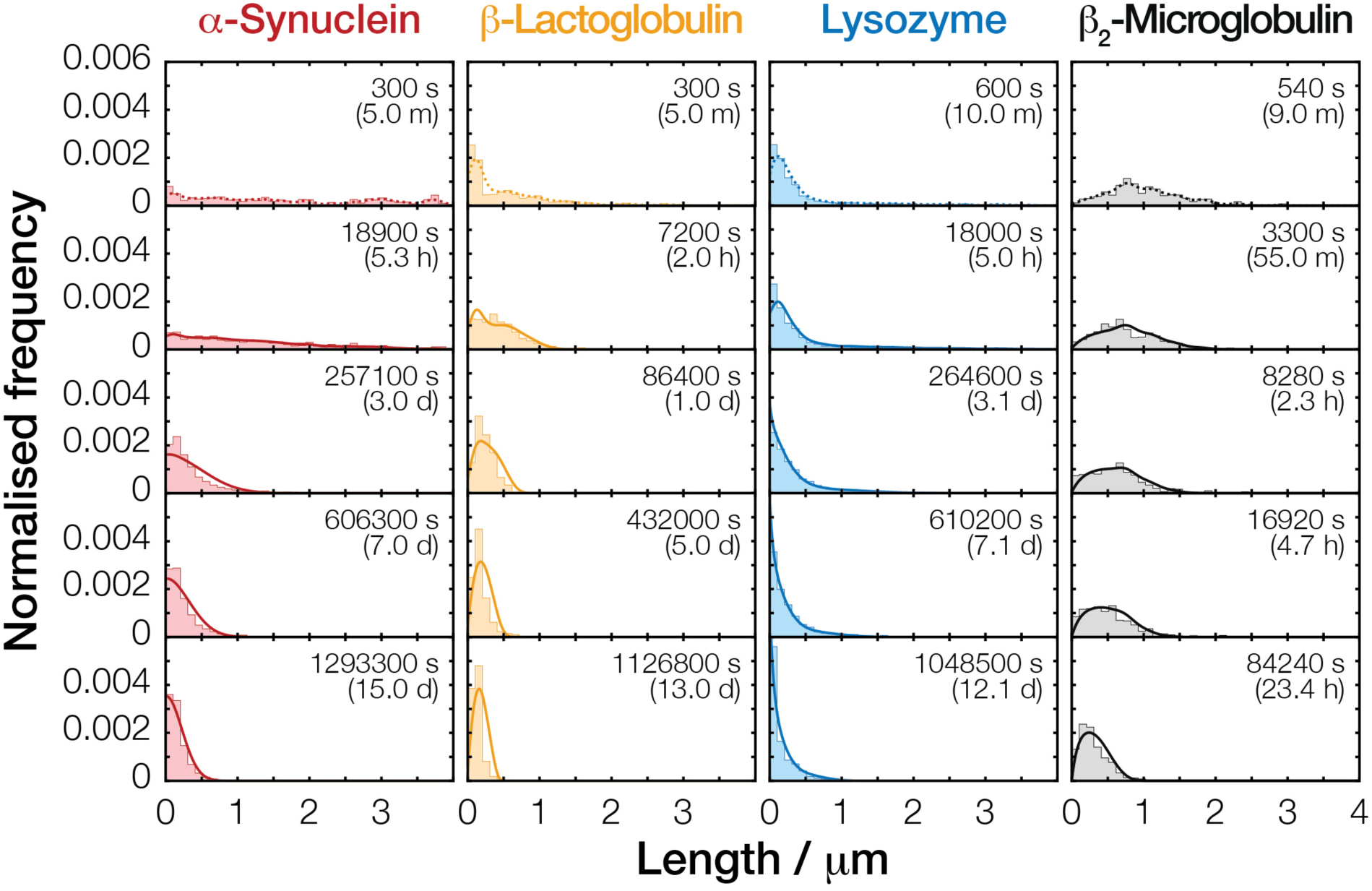
Validation of the division parameters α, γ and κ_0_ and their predictive power. Full direct simulation of fibril fragmentation processes using α, γ and κ_0_ determined from the image data. For each fibril type, the initial normalised frequency distribution (dashed lines in top row) were used directly as the initial state for the simulations. The resulting simulated evolution of length distributions solely based on the calculated α and γ values and estimated shapes κ_0_ (see Materials and methods) are compared with the experimental data show as histograms.

## DISCUSSION

Understanding of the properties that underline the biological activities of amyloid nano-structures, such as their cytotoxic and infectious potentials, is crucial for the understanding of why some amyloid is associated with devastating human diseases. The division of amyloid fibrils, for example through fibril fragmentation by mechanical perturbation ^22, 35^, enzymatic action ^51-52^ or other cellular or environmental perturbations, is a key step in their life-cycle that results in the exponential growth in the number of amyloid particles. Simultaneously, daughter particles resulting from the division of parent fibrils cause a reduction in the overall size distribution as division proceeds. These two consequences of division are undoubtedly linked to the enhancement of the cytotoxic and infectious potentials of disease-associated amyloid ^20-21^. The amyloid fibrils’ resistance towards division, i.e. the stability of the amyloid fibrils towards division, therefore, rationalise the two fundamental requirements for pathogenicity associated with amyloid. Akin to uncontrolled division of cells or any pathogenic microorganisms, the division step in the amyloid life cycle (**Fig. 1**) could be a key determinant in their overall potential to be associated with properties in the amyloid and prion associated pathology.

Here, we have developed a theory, as well as an experimental approach utilising our theoretical insights to resolve the amyloid fibrils’ dynamic stability towards division. These represent a step change in how we can study amyloid fibril division processes such as in fibril fragmentation and prion propagation, essentially the replication step in the amyloid lifecycle. It also allows the direct comparison between amyloid particles of different molecular types and quantifies the difference in division and stability between those that are and are not disease associated. Specifically, we have applied our theoretical results to the comparison of a diverse set of amyloid assemblies consisting of human α-Syn (a neurodegenerative disease-associated amyloid, sample formed under physiological solution conditions), human β_2_m (a systemic amyloidosis disease-associated amyloid, sample formed under acidic pH, data from Xue and Radford 2013 ^35^), bovine β-Lac and hen egg white Lyz (later two cases are both biophysical model systems not directly related to human disease but converted to amyloid when subjected to heating in acidic pH). By analysing and comparing their division behaviour fully and uniquely described by the triplet of parameters: *α* (magnitude of the division rate constant), *γ* (fibril length dependence of the division rate constant), and *κ*_*0*_ (probability of division at any given position along a fibril) under identical mechanical perturbation for long timescales using our approach, we show a remarkable difference in the stability of these different amyloid assemblies relative to each other and how they divide (summarised in **Fig. 8**). Interestingly, for the four fibril types we included here, considering the division rate constant B with their cross-sectional area, the disease associated human α-Syn fibrils demonstrate lowest overall stability towards division followed by Lyz, human β_2_m and finally β-Lac particles that are most stable towards division (**Fig 8.** last row). Based on the comparison of the *α* and *γ* parameters that together describe the division rates B(x), the likelihood that small α-Syn particles (<100 nm long) will divide is similar to that of Lyz particles of identically length despite having more than double the mean width (and thus around 4 times bigger cross-sectional area, **Table 1** and **Fig. 8**). More importantly, the division of α-Syn particles also results in a larger relative concentration of small particles compared to β_2_m and β-Lac. These results show that human α-Syn amyloid fibrils are relative unstable assemblies capable of a more rapid shedding of small particles that could well possess enhanced cytotoxic and infectious potentials ^53^ through division compared with the other fibril types investigated here. Thus, our results also directly suggest a testable causality link between the low stability of α-Syn fibrils towards division and recent observations that human α-Syn may behave in a prion-like manner in cell to cell propagation and their cytotoxicity ^54^.

**Figure 8.**
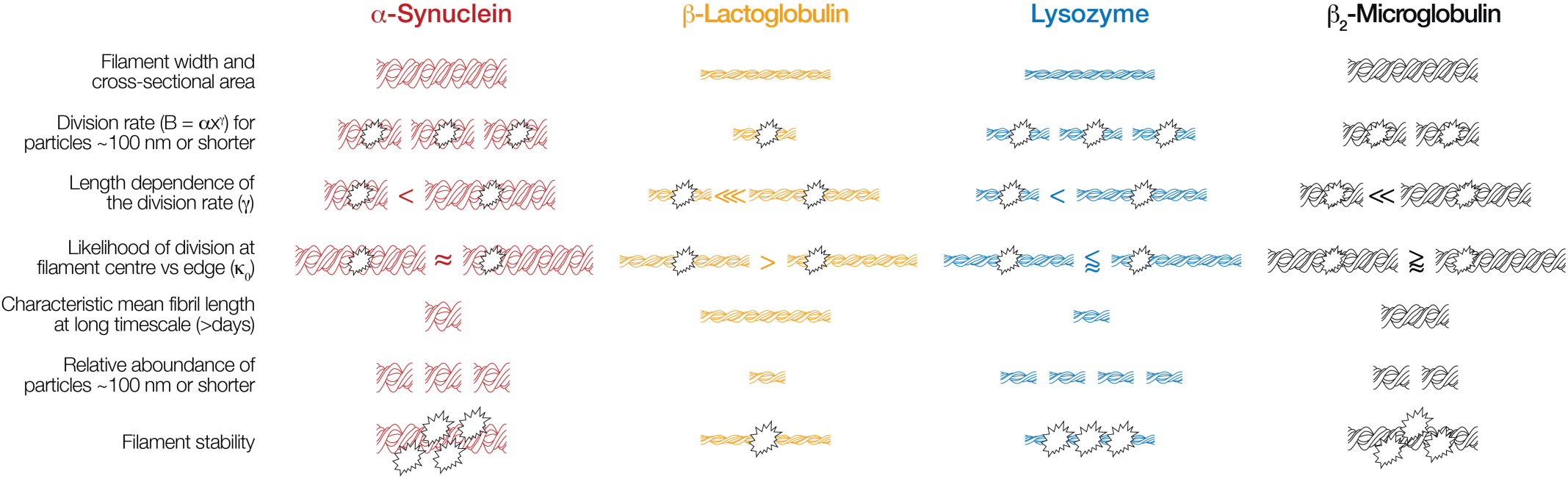
Schematic summary of the fibril division properties and their consequences compared between each of the fibril types. Comparison of the fibril division profiles reveal differences in the dynamical stability towards breakage for the four different types of amyloid fibrils, and suggest that disease-related amyloid have lowered stability towards breakage and increased likelihood of shedding smaller particles compared to amyloid not related to disease. In the illustrations, the fibril width, number and number of breakage symbols are not to scale and denote the relative rankings for the different properties.

Since the division of amyloid fibrils is an integral part in the propagation of the amyloid conformation (**Fig. 1**), the nanoscale materials properties of amyloid underpin processes which drive the proliferation of amyloid, as well as their varied roles in biology. Therefore, it is important to appreciate the suprastructural properties of amyloid (e.g. clustering, bundling, twist, stiffness, width distribution, orientation distribution, and length distribution etc) at mesoscopic (nanometre to micrometre) length scales, as these properties will influence how individual amyloid fibrils divide. Our data show that despite all amyloid consisting of a cross-beta core structure, their ability to resist division through fragmentation promoted by mechanical perturbation varies strongly between fibril types. Since the stability of amyloid fibrils towards division will depend on their suprastructural properties, which in turn depends on their precise structure at atomic level, mesoscopic level structural properties may well be the missing link between amyloid structure and the varied biological effects and consequences that different amyloid types evoke under different conditions. Thus, it should be possible to generate a structure activity relationship (SAR) correlating the suprastrucutral properties of amyloid, their ability to divide, and their cytotoxic and/or infectious potentials. Understanding this SAR for amyloid assemblies could lead to designer bio-safe polymers with tuned mechanical and nanomaterials properties as well as rationalise the disease associated properties of amyloid structures.

Analogous to the diverse response of soluble folded proteins towards unfolding by chemical denaturants, thermal melting and mechanical force etc., the stability of amyloid fibrils could also vary depending on the nature of the perturbation. Indeed, amyloid fibrils may break down in the presence of chemical, thermal or enzymatic action ^15, 51-52, 55-57^, and their relative resistance or stability towards different stresses, including those associated with physiological changes involved in human disorders, is not known. In particular, understanding how enzymatic action by molecular chaperones such as Hsp104 or ClpB promote amyloid division, degradation and/or propagation of amyloid conformation ^58-59^ in relevant cases may be key in resolving the complex behaviour of the amyloid lifecycle in a biological context. In summary, the combined theoretical and experimental work we report here will enable the characterisation and comparison of the amyloid division processes and the relative stabilities of amyloid assemblies. Both properties are fundamental in understanding the lifecycle of disease-associated amyloid as well as the normal roles of functional amyloid in biology.

## MATERIALS AND METHODS

### Preparation of protein monomers

Hen egg white Lyz and bovine β-Lac proteins were both purchased from Sigma-Aldrich and used with no further purification. Production and purification of human α-Syn monomers was achieved according to the method of Cappai et al ^60^, with the addition of a stepped ammonium sulphate precipitation (30% to 50%) step prior to anion exchange chromatography. The protein was buffer exchanged using PD10 desalting column (GE Healthcare) prior to loading onto the anion exchange resin.

### *In vitro* formation of amyloid fibril samples

The conversion of Lyz and β-Lac to amyloid fibres was achieved under acidic and heated conditions. Both proteins were dissolved in 10 mM HCl to a concentration of 15mg/ml and then incubated for 4 hr at 25 °C. The resulting solutions were filtered through a 0.2 *µ*m syringe filter and diluted to a concentration of 10mg/ml (Lyz = 699 *µ*M and β-Lac = 547 *µ*M). 500 *µ*l aliquots were then heated without agitation for differing periods of time, with Lyz heated at 60 °C for 2 days and β-Lac heated at 90 °C for 5 hr. α-Syn fibrils were formed by buffer exchange of purified monomers into fibril forming buffer (20mM Sodium phosphate, pH7.5) using a PD-10 column (GE Healthcare). The resulting α-Syn solution was passed through a 0.2 *µ*m syringe filter. Protein concentration was subsequently determined via absorbance at 280nm, and the sample solution were diluted to 300 *µ*M and incubated at 37 °C in a shaking incubator with agitation set at 200 rpm for at least two weeks.

### Controlled fibril fragmentation through mechanical perturbation

Parent fibril solutions were diluted to 120 *µ*M using the appropriate fibril forming buffer for each protein in a snap cap vial containing an 8 x 3 mm PTFE coated magnetic stirrer bar and then subjected to stirring at 1000 rpm on an IKA squid stirrer plate with digital speed display. At appropriate time points, small aliquots of the fibril samples were removed, diluted with fibril forming buffer (deposition concentration for α-synuclein is 0.48 *µ*M, β-lactoglobulin is 0.6 *µ*M and Lyz is 6 *µ*M), and 20 *µ*l were immediately taken and incubated for 5 min on freshly cleaved mica surfaces (Agar Scientific F7013). The mica surfaces were subsequently washed with 1 ml of syringe filtered (0.2 *µ*m) mQ H_2_O and dried under a gentle stream of N_2_(g).

### Determination of residual monomer concentration

Residual monomer concentration for each fragmentation sample were measured using SDS-PAGE after centrifugation (75000 rpm, 15 min) with 100 *µ*l of the 120 *µ*M fragmentation reaction and 100 *µ*l of 120 *µ*M non-fragmented parent fibrils samples. The top 10*µ*l of the solutions were then removed and treated with 4x loading dye and boiled at 95 °C for 5 min (Lyz samples were heated to 65 °C and beta-mercaptoethanol was not added due to decomposition of samples). The samples were then run against a serial dilution of monomeric protein standards on either a Tris-Tricine gel or a 15% Tris-Glycine gel at 180V and subsequently stained with Coomassie blue. Analysis of the protein bands were carried out by densitometry for comparison of bands to the serial dilution bands.

### AFM imaging and image analysis

The fibril samples were imaged on a Bruker Multimode 8 scanning probe microscope with a Nanoscope V controller, using the ScanAsyst peak-force tapping imaging mode. Bruker ScanAsyst probes (Silicone nitride tip with tip height = 2.5-8 μm, nominal tip radius = 2 nm, nominal spring constant 0.4 N/m and nominal resonant frequency 70 kHz) were used throughout. Multiple 20 *µ*m x 20 *µ*m areas of the surface were scanned at a resolution of 2048 x 2048 pixels. The images were then processed and flattened using Bruker Nanoscope Analysis software to remove tilt and bow. The images were then imported into Matlab, length of individual fibril particles was measured and the sample length and height distributions were obtained as previously described ^47, 61^.

### Data analysis of fibril division properties

The normalised length distribution of the fibril samples measured by AFM at time *t, f* (*t, x*), is linked to the concentration of fibrils solution *u* (*t, x*) in Eq. (1) by the relation in Eq. (SI.11). Mean lengths for each time point *μ*(*t*) were calculated from the experimental *f* (*t, x*) distributions and Eq. (2) and (4) where used to first extract *γ* from the datasets. Because some unknown number of experimentally measured length distributions at early time points in the experiments may be at considerably distance from the self-similar distribution shape and the stability line (i.e. where Eq. (2) does not apply), we fit the following equation Eq. (4) to the average length as function of time data instead of Eq. (2) directly in order to estimate the number of experimental time points consistent with the self-similar distribution shape objectively without human input:

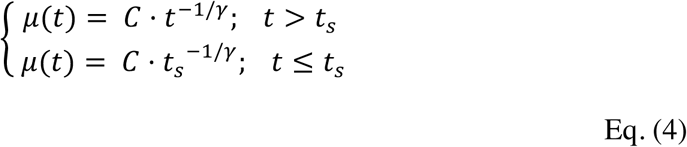

Eq. (4) was fit to the average length *μ*(*t*) as function of time *t*, with *C* and *t*_*s*_ as parameters individual to each experimental dataset and *γ* as a global parameter for datasets from the same fibril type. Subsequently, the *g*(*x*_*g*_) and *α* values were calculated with Eq. (3) and (SI.21), respectively, both using *γ* calculated above and experimental normalised length distributions *f*(*t, x*) where *t* > *t*_*s*_. For both the *g*(*x*_*g*_) distributions and *α* values, averages were obtained for each fibril type. The self-similar distribution shapes *g*(*x*_*g*_) were used to calculate length distribution at any time using the reverse of Eq. (3). The *α* and *γ* values were used to calculate the division rate constant *B*(*x*) = *α*_0_(*αx*)^*γ*^ for fibrils of any length *x*. Supplementary information section contains further information on the mathematical considerations of our division model.

### Direct numerical simulation of fibril division processes

To validate the *α* and *γ* values obtained from our analysis, direct numerical simulations to calculate the time evolution of the fibril length distributions were carried out by numerically solving the full ODE system describing the master equation mostly as described in Xue and Radford 2013 ^35^ but with few modifications. Firstly, numerical integrations of the master equation were solved for fibril species containing up to 30,000 instead of 20,000 monomeric units in order to retain concentration errors introduced by numerical inaccuracy and truncation of larger species to <1%. Secondly, the number of division sites were assumed to be equal to the number of monomers-1 and the unit used for the length of fibrils were interconverted in the simulations from nanometres (*x* in [nm] units) to the number of monomers (*i* number of monomers) using the numbers of monomers per nm length unit *N*_*l*_ ^35^ as conversion factor. Subsequently, assuming that division sites along the fibrils operate independently, the microscopic rate constant on per division site basis is *B(i)κ*_*0*_ divided by the number of monomers-1. Thirdly, as *g*(*x*_*g*_) shape for Lyz and *α*-Syn fibril divisions suggest a *κ*_*0*_ function that result in similar division rates in the fibril centre and fibril edge, simulations for these two fibril types were carried out using Eq. S6 in Xue and Radford 2013 instead of Eq. S8. Finally, the experimental distribution at the first time-points (including all the experimental noise) were directly used as the initial distribution (dashed lines in **Fig. 6**) instead of parameterised distributions ^61^ in the simulations to test the fact that our model has shown that the self-similar distribution shape will be reached independently of the initial length distribution.

## ACKNOWLEDGEMENTS

We thank the members of the Xue group, and the Kent Fungal Group for helpful comments throughout the preparation of this manuscript, and Ian Brown for technical support. We also thank Sheena Radford for insightful discussions, as well as help and support on the reanalysis of β_2_-microglobuling fragmentation dataset. This work was supported by funding from Inria, and the Biotechnology and Biological Sciences Research Council (BBSRC), UK grants BB/J008001/1 as well as BB/F016719/1 (D.M.B).

