## Supplementary Information for "The division of amyloid fibrils – Systematic comparison of fibril fragmentation stability by linking theory with experiments"

### Supporting Information

*David M. Beal[1], Magali Tournus[2], Ricardo Marchante[1,6], Tracey Purton[1], David P. Smith[3], Mick F. Tuite[1], Marie Doumic[4,5], Wei-Feng Xue[1,4]**

[1]Kent Fungal Group, School of Biosciences, University of Kent, CT2 7NJ, Canterbury, UK, [2]Centrale Marseille, I2M, UMR 7373, CNRS, Aix-Marseille univ., Marseille, 13453, France [3]Biomolecular Research Centre, Sheffield Hallam University, Sheffield, UK [4]INRIA Rocquencourt, équipe-projet BANG, domaine de Voluceau, BP 105, 78153 Rocquencourt, France [5]Wolfgang Pauli Institute, University of Vienna, Vienna, Austria [6]Current address: Institute for Genetics and CECAD Research Center, University of Cologne, Joseph-Stelzmann Str. 26, 50931 Cologne, Germany

Key words: amyloid and prions / atomic force microscopy / fragmentation / breakage / dynamic stability / self-similar length distribution / mathematical model

**SUPPORTING INFORMATION**

**The self-similar division equation**

We first explain here the origins of Eq. (1) in the Main Text as well as the assumptions associated with this equation. Let us denote the fibril length distribution *u(t, x)* as the particle concentration of fibrils of length x > 0 at time t, *B(x)* $\geqslant$ *0* as the division rate constant for fibrils of length *x* (assumed to be independent of time), and *κ(y, dx)* the probability that a dividing fibril of length *y* gives rise to two fibrils of size *x* and *y* − *x* (Fig. 1b). The *κ*, often called fragmentation kernel, is nonnegative and satisfies the following properties:

$$\int_{0}^{y} \kappa(y, dx) = 1, \qquad \kappa(y, x > y) = 0, \qquad \kappa(y, x) = \kappa(y, y - x)$$

Eq. (SI.1)

The last property above is a symmetry property linked to the assumption that we consider only division into two daughter fibrils for each microscopic step, and the fibrils are isotropic along the axis of the filament so the division rate only depends on the length of the resulting two fibrils (**Fig. 1c** and **1d**). The first two properties of Eq. (SI.1) express that *κ(y, dx)* is a normalised probability density function, and that daughter fibrils post-division are always shorter than their mother fibril. The time dependent concentration of fibrils *u(t, x)* then satisfies the following equation:

$$\frac{\partial}{\partial t}u(t,x) = -B(x)u(t,x) + 2\int_{y=x}^{\infty} \kappa(y,x)B(y)u(t,y)dy, \qquad u(0,x) = u_0(x)$$

Eq. (SI.2)

where $u_0(x)$ is the initial length distribution of fibrils. Equation (SI.2) is the continuous division equation, which describes the evolution $\frac{\partial}{\partial t}u(t,x)$ of the fibril particle concentrations in the fibril length distribution $u(t, x)$ with respect to time $t$. It states that fibrils of a given length $x$ in the sample distribution will be consumed with a rate $B(x)$ when they divide into smaller daughter fibrils, and that fibrils of the same length $x$ may also appear in the sample distribution each time a fibril of size $y > x$ divides into two fibrils of size $x$ and $y - x$. Let us denote the total initial mass of fibrils as $\rho = \int_0^\infty x u_0(x) dx$. Since the mass is conserved through time: $\int_0^\infty x u(t, x) dx = \rho$. We also assume, in line with previous theoretical [33] and experimental results [35], that the division rate constant is given by a power law:

$$B(x) = \alpha_0 (\alpha x)^\gamma, \qquad \alpha > 0, \qquad \gamma > 0$$

Eq. (SI.3)

and that the site where a fragmenting fibril of size $y$ breaks down only depend on the relative position of its site along the mother fibril, defined by the ratio $x/y$ where $x$ is the length of one of the two daughter fibrils. This property is called a "self-similar" division and is translated mathematically with fragmentation kernel $\kappa$ as the following:

$$\kappa(y, x) := \frac{1}{y} \kappa_0 \left( \frac{x}{y} \right)$$

Eq. (SI.4)

where the properties described by Eq. (SI.1), when transferred to the probability density $\kappa_0$, and with $z = \left(\frac{x}{y}\right)$, satisfies the following:

$$\int_0^1 \kappa_0(z) dz = 1, \qquad \kappa_0(z > 1) = 0, \qquad \kappa_0(z) = \kappa_0(1 - z)$$

Eq. (SI.5)

Two important examples may be viewed as special cases of self-similar fragmentation kernels above. The first one is the case of division of uniform probability: the mother fibril can break at any site along its length with an equal probability, so that $\kappa_0 \left( \frac{x}{y} \in (0,1) \right) = 1$. The second special division case is sometimes referred to as the "equal mitosis case" from its roots in describing cellular divisions, where the mother fibril divides exactly at the middle, so that we have a Dirac delta function at $\kappa_0\left(\frac{1}{2}\right)$: $\kappa_0\left(\frac{x}{y}\right) = \delta_{\frac{x}{y}=\frac{1}{2}}$. Using all of the properties and assumptions above, the continuous division equation Eq. (SI.2) then becomes:

$$\frac{\partial}{\partial t}u(t,x) = -\alpha_0(\alpha x)^{\gamma}u(t,x) + 2\int_{y=x}^{\infty}\frac{1}{y}\kappa_0\left(\frac{x}{y}\right)\alpha y^{\gamma}u(t,y)dy$$

Eq. (SI.6)

which is equation Eq. (1) in the Main text.

#### Long-time behaviour of the continuous division equation

For our continuous division equation Eq. (1) and (SI.6), it has been proven in [49] that for long times, there exists a unique probability density function *g* and a constant $C_g > 0$ such that:

$$u(t,x) \underset{t\to\infty}{\longrightarrow} C_g t^{\frac{2}{\gamma}} g(x_g), \qquad x_g = x t^{\frac{1}{\gamma}}$$

Eq. (SI.7)

The constant $C_g$ is introduced to ensure mass conservation, which holds for any time *t*. Eq. (SI.7) means that for large times, the probability density *u* tends towards a specific distribution shape *g* after variable rescaling. Moreover, the function *g* is defined as the unique solution to the following equation:

$$x_g\frac{dg(x_g)}{dx_g} + (2 + \alpha\gamma {x_g}^{\gamma})g(x_g) = 2\alpha\gamma\int_{y_g=x_g}^{\infty}\frac{1}{y_g}\kappa_0\left(\frac{x_g}{y_g}\right)y_g^{\gamma}g(y_g)dy_g, \qquad \int_0^{\infty}g(y_g)dy_g = 1$$

Eq. (SI.8)

We can then compute the constant $C_g$ as the following:

$$\int_0^{\infty}xu(t,x)dx = \rho = C_g\int_0^{\infty}t^{\frac{2}{\gamma}}xg\left(xt^{\frac{1}{\gamma}}\right)dx = C_g\int_0^{\infty}x_g g(x_g)dx_g \quad\Rightarrow\quad C_g = \frac{\rho}{\int_0^{\infty}x_g g(x_g)dx_g}$$

Eq. (SI.9)

We then relate these results to our experimental measurements. First, since we measure at successive time points small aliquots taken from the fibril samples, these samplings may be viewed as measurements of the length distribution of the fibril sample at time points *t*. We also do not measure directly *u(t, x)*, since the total number of fibrils is not known *a priori* for each time point. Instead, we measure the normalised length distribution *f(t, x)* as described below. Using Eq. (SI.7-9), we then have the following:

$$\int_0^\infty u(t,x)dx \underset{t\to\infty}{\longrightarrow} C_g \int_0^\infty t^{\frac{2}{\gamma}} g\left(xt^{\frac{1}{\gamma}}\right) dx = C_g t^{\frac{1}{\gamma}} \int_0^\infty g(x_g)dx_g = C_g t^{\frac{1}{\gamma}}$$

Eq. (SI.10)

We can define $f(t,x)$ as the normalised fibril length distribution that can be assessed using the experimental image data:

$$f(t,x) = \frac{u(t,x)}{\int_0^\infty u(t,x)dx}$$

Eq. (SI.11)

Using this definition of *f(t, x)* from, we then have:

$$f(t,x) \underset{t\to\infty}{\longrightarrow} \frac{C_g t^{\frac{2}{\gamma}} g(x_g)}{C_g t^{\frac{1}{\gamma}}} = t^{\frac{1}{\gamma}} g(x_g), \qquad x_g = xt^{\frac{1}{\gamma}}$$

Eq. (SI.12)

which is equation Eq. (3) of the main text. Next, defining the average length of fibrils *μ(t)* as the experimentally tractable time-dependent mean length of the fibril length distribution defined as:

$$\mu(t) = \int_0^{\infty} x \cdot f(t, x) dx$$

Eq. (SI.13)

We have the following relationship:

$$\mu(t) := \int_0^{\infty} xf(t,x)dx \underset{t\to\infty}{\longrightarrow} \int_0^{\infty} xt^{\frac{1}{\gamma}} g\left(xt^{\frac{1}{\gamma}}\right) dx = t^{-\frac{1}{\gamma}} \int_0^{\infty} x_g g(x_g) d\square_g = Ct^{-\frac{1}{\gamma}}, \qquad C = \int_0^{\infty} x_g g(x_g) dx_g$$

Eq. (SI.14)

which is the relationship between the average length of fibrils and time *t* in equation Eq. (2) of the main text.

**Estimating the division parameters α and γ**

We first estimate $\gamma$ by fitting a modified version of Eq. (2) to the average lengths $\mu(t)$ estimated from the experimentally observed fibril length distributions for sufficiently long times (see Eq. (7) in Materials and Methods). Then, we estimate $\alpha$ from $\gamma$ and *g* using Eq. (SI.8). Integration of Eq. (SI.8) yields:

$$\int_0^{\infty} x_g \frac{dg(x_g)}{dx_g} dx_g + \int_0^{\infty} 2g(x_g) dx_g + \alpha\gamma \int_0^{\infty} x_g{}^{\gamma} g(x_g) dx_g$$

$$= 2\alpha\gamma \int_0^{\infty} \int_{y_g = x_g}^{\infty} \frac{1}{y_g} \kappa_0 \left(\frac{x_g}{y_g}\right) y_g^{\gamma} g(y_g) dy_g \, dx_g$$

Eq. (SI.15)

We can integrate Eq. (SI.15) by parts the first term, and we use Fubini's theorem to invert the integral order in the last term:

$$-\int_0^\infty g(x_g)dx_g + \int_0^\infty 2g(x_g)dx_g + \alpha\gamma\int_0^\infty {x_g}^\gamma g(x_g)dx_g$$

$$= 2\alpha\gamma\int_0^\infty \int_{x_g=0}^{y_g} \frac{1}{y_g}\kappa_0\left(\frac{x_g}{y_g}\right) y_g^\gamma g(y_g)dx_g\, dy_g$$

Eq. (SI.16)

We then use the fact that $g$ is normalised, $\int_0^\infty g(y_g)dy_g = 1$, and change the variable $x_g$ to $z = \left(\frac{x_g}{y_g}\right)$ to obtain:

$$1 + \alpha\gamma\int_0^\infty {x_g}^\gamma g(x_g)dx_g = 2\alpha\gamma\int_0^\infty \int_{z=0}^{1} \kappa_0(z) y_g^\gamma g(y_g)dz\, dy_g$$

Eq. (SI.17)

Using the property $\int_0^1 \kappa_0(z)dz = 1$ from Eq. (SI.1), we obtain:

$$1 = \alpha\gamma\int_0^\infty {x_g}^\gamma g(x_g)dx_g$$

Eq. (SI.18)

To relate $\alpha$ directly to the experimentally characterised *f(t,x)* rather than on $g$, we multiply the equation Eq. (SI.12), i.e. Eq. (3) of the main text, by $x^\gamma$ and integrate it to obtain the following:

$$\int_0^\infty x^\gamma f(t,x)dx \underset{t\to\infty}{\longrightarrow} \int_0^\infty x^\gamma t^{\frac{1}{\gamma}} g\left(xt^{\frac{1}{\gamma}}\right)dx = \int_0^\infty x_g^\gamma t^{-1} g(x_g)dx_g$$

Eq. (SI.19)

Rearranging Eq. (SI.18) and using Eq. (SI.19), we obtain:

$$\alpha = \frac{1}{\gamma}\frac{1}{\int_0^\infty {x_g}^\gamma g(x_g)dx_g} \underset{t\to\infty}{\longrightarrow} \frac{1}{\gamma}\frac{t^{-1}}{\int_0^\infty x^\gamma f(t,x)dx}$$

Eq. (SI.20)

Therefore, we get the following relationship:

$$\alpha \approx \frac{1}{\gamma} \frac{t^{-1}}{\int_0^\infty x^\gamma f(t,x)dx}, \qquad t \gg t_0$$

Eq. (21)

which is used to estimate $\alpha$ from experimental data. For more details, we also refer the interested reader to Doumic et al. [50], and more specifically to Lemma 1 and Eq. (3.3) in this reference.

#### Supporting figures and table

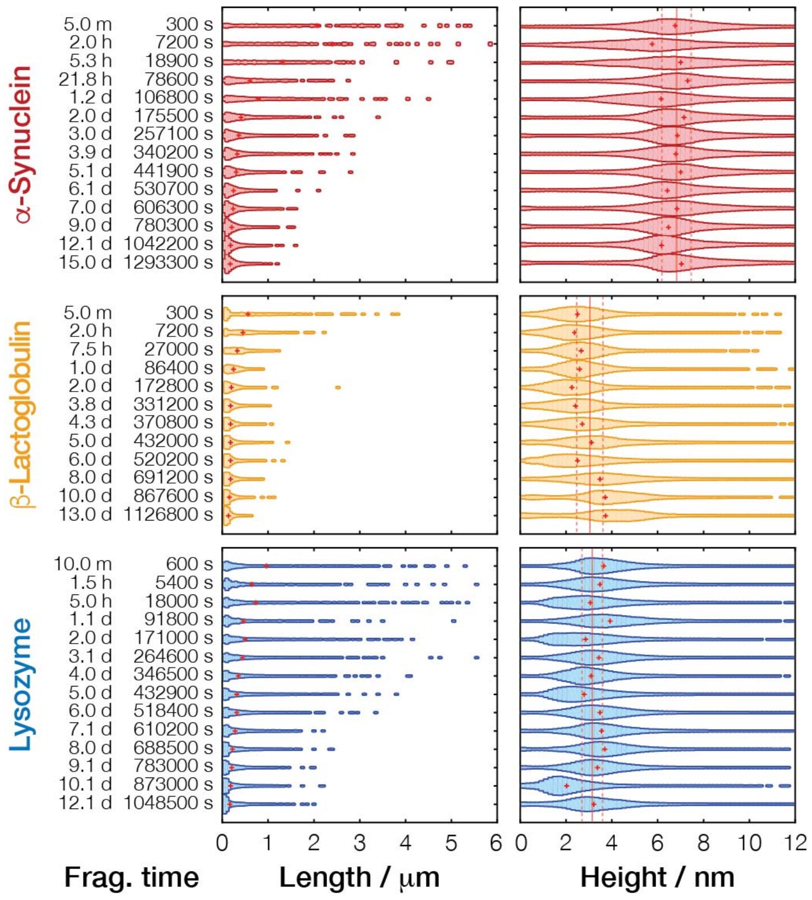


**Supporting Figure S1. Fibril length and height distributions extracted from AFM images for fibrils undergoing fragmentation by mechanical stirring.** Typical experimental time course with normalized length (left plot of each sample) and height (right plot of each sample) distributions of fibril particles shown as violin plots. The width of the horizontal bars corresponds to the normalised frequencies observed at the length or height indicated by the x-axes. The bars for all samples are shown using the same length and height frequency scales, respectively, to facilitate comparison. The red crosses indicate mean values at each time point and the solid and dashed red lines for height plots indicate mean and standard deviation of all time points taken together, respectively.

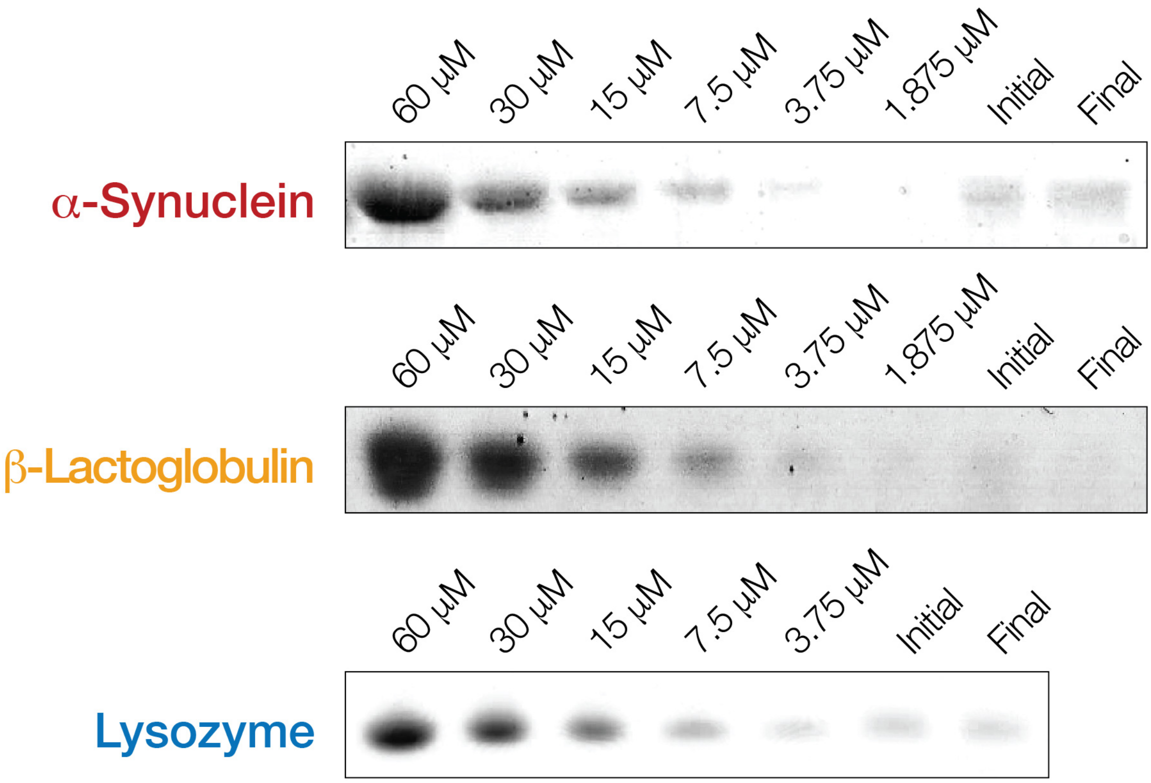


**Supporting Figure S2: Residual monomer assay before and after fibril fragmentation time courses.** For each fibril type, protein content in the non-pellatable fractions of the initial sample before and Final sample after extended mechanical perturbation were visualised on SDS-PAGE gels together with loading standards of known protein concentrations. The difference in residual monomer concentration (difference between bands in the Initial and Final lanes) were less than 5 % in all cases.

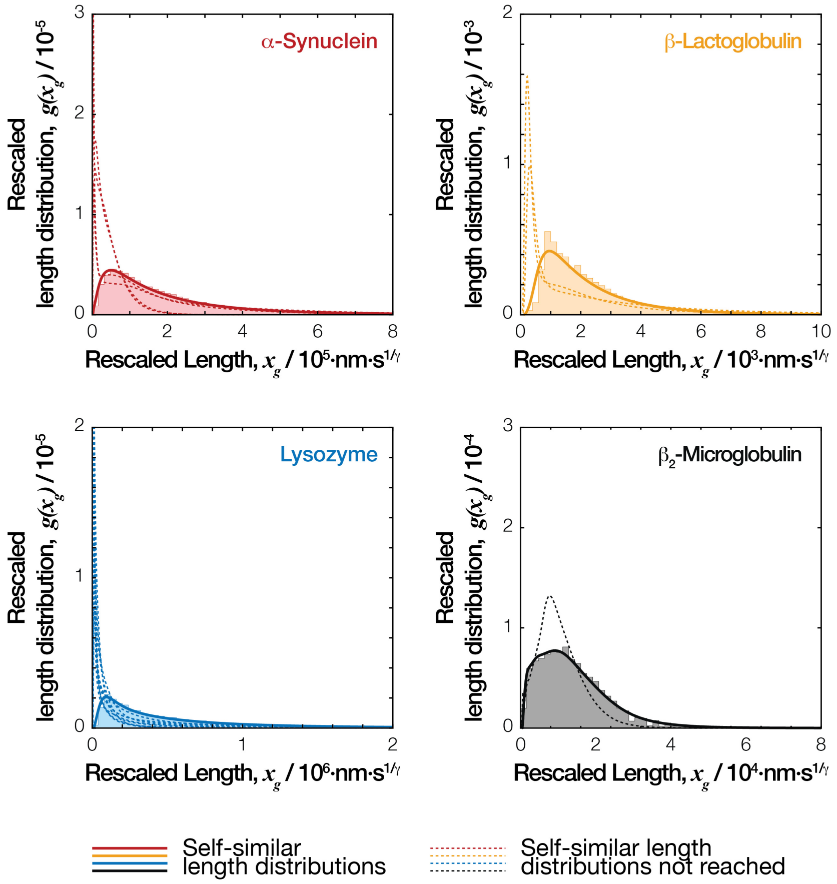


**Supporting Figure S3: The self-similar length distribution shape can be obtained from rescaling and averaging of the experimental normalised length distributions.** The resealed length distributions $g(x_g)$ calculated with Eq. (5) are shown for each fibril type. For each fibril type analysed, the histograms and bold solid lines are the average of length distributions obtained from AFM imaging analysis that have reached the self-similar length distribution shapes, i.e. distributions at the time points consistent with Eq. (2) in the portion of the experiments represented by the solid lines in **Fig. 5**. for each fibril type. The dashed lines represent distributions from early experimental time-points where self-similarity has not been reached, demonstrating the large deviations from the self-similar distribution shape represented by the bold lines. The lines represent distributions calculated using the kernel density method to reduce clutter and facilitate visualisation and comparison.

**Supporting Table S1:** Sample, AFM imaging and quantitative image analysis statistics.

| ***Samples*** | | ***AFM Imaging*** | | | ***Quantitative image Analysis*** | | | |
|---|---|---|---|---|---|---|---|---|
| *Protein* | *Fragmentation Time /s* | *Number of Images* | *Image size / pixels*† | *Scan size / μm*† | *Mean Particle Length / nm* | *Number of Fibril Particles*◊ | *Mean Particle Height / nm* | *Number of Pixels*° |
| ***α-Syn*** | 300 | 6 | 2048 | 20 | 2075.8 | 653 | 6.8 | 49079 |
| | 7200 | 5 | 2048 | 20 | 2402.9 | 424 | 5.8 | 32066 |
| | 18900 | 3 | 2048 | 20 | 1311.6 | 679 | 7.0 | 43259 |
| | 78600 | 2 | 2048 | 20 | 598.9 | 1350 | 7.3 | 53512 |
| | 106800 | 2 | 2048 | 20 | 786.8 | 1512 | 6.1 | 63474 |
| | 175500 | 2 | 2048 | 20 | 406.2 | 3086 | 7.1 | 92632 |
| | 257100 | 2 | 2048 | 20 | 360.1 | 4087 | 6.8 | 110048 |
| | 340200 | 2 | 2048 | 20 | 330.6 | 4257 | 6.8 | 110024 |
| | 441900 | 2 | 2048 | 20 | 301.8 | 2723 | 7.0 | 67278 |
| | 530700 | 2 | 2048 | 20 | 244.7 | 5064 | 6.4 | 110431 |
| | 606300 | 2 | 2048 | 20 | 241.4 | 6104 | 6.8 | 129898 |
| | 780300 | 2 | 2048 | 20 | 208.9 | 6707 | 6.5 | 129581 |
| | 1042200 | 2 | 2048 | 20 | 185.6 | 6412 | 6.1 | 113509 |
| | 1293300 | 2 | 2048 | 20 | 173.6 | 5346 | 7.0 | 90832 |
| | 300 | 5 | 2048 | 20 | 2183.7 | 507 | 7.5 | 37725 |
| | 5400 | 5 | 2048 | 20 | 1756.2 | 394 | 7.0 | 24323 |
| | 18000 | 2 | 2048 | 20 | 1303.7 | 635 | 7.8 | 36938 |
| | 91800 | 2 | 2048 | 20 | 620.7 | 2293 | 7.2 | 87165 |
| | 107400 | 2 | 2048 | 20 | 568.5 | 2304 | 7.4 | 80443 |
| | 195900 | 2 | 2048 | 20 | 439.7 | 2416 | 7.2 | 77497 |
| | 259200 | 2 | 2048 | 20 | 390.5 | 3264 | 7.1 | 93323 |
| | 430500 | 2 | 2048 | 20 | 276.7 | 4807 | 6.9 | 114822 |
| | 610200 | 2 | 2048 | 20 | 224.3 | 6253 | 7.1 | 128690 |
| | 691200 | 2 | 2048 | 20 | 205.2 | 7541 | 6.8 | 145181 |
| | 766200 | 2 | 2048 | 20 | 194.0 | 5705 | 6.4 | 106589 |
| | 863700 | 2 | 2048 | 20 | 181.7 | 8086 | 6.5 | 143206 |
| | 1119600 | 2 | 2048 | 20 | 172.0 | 9532 | 6.4 | 161606 |
| | 1219800 | 2 | 2048 | 20 | 163.6 | 9470 | 6.3 | 154357 |
| | 1380900 | 2 | 2048 | 20 | 158.3 | 8811 | 6.8 | 139488 |
| ***β-Lac*** | 300 | 2 | 2048 | 20 | 560.2 | 496 | 2.5 | 28968 |
| | 7200 | 2 | 2048 | 20 | 451.8 | 854 | 2.4 | 40389 |
| | 27000 | 1 | 2048 | 20 | 329.7 | 751 | 2.7 | 26125 |
| | 86400 | 1 | 2048 | 20 | 244.7 | 1023 | 2.6 | 26678 |
| | 172800 | 1 | 2048 | 20 | 200.8 | 1482 | 2.3 | 32010 |
| | 331200 | 2 | 2048 | 20 | 183.0 | 2637 | 2.4 | 52129 |
| | 370800 | 1 | 2048 | 20 | 186.0 | 1112 | 2.7 | 22330 |
| | 432000 | 2 | 2048 | 20 | 185.2 | 2180 | 3.1 | 43588 |
| | 520200 | 2 | 2048 | 20 | 181.1 | 2190 | 2.5 | 42867 |

| | | | | | | | | |
|---|---|---|---|---|---|---|---|---|
| | 691200 | 2 | 2048 | 20 | 181.0 | 1945 | 3.5 | 38036 |
| | 867600 | 1 | 2048 | 20 | 163.2 | 922 | 3.7 | 16349 |
| | 1126800 | 1 | 2048 | 20 | 134.2 | 1138 | 3.7 | 16819 |
| | 1800 | 3 | 2048 | 20 | 680.8 | 898 | 2.0 | 63514 |
| | 3600 | 2 | 2048 | 20 | 495.8 | 977 | 2.9 | 50594 |
| | 87588 | 1 | 2048 | 20 | 304.6 | 2649 | 3.1 | 85313 |
| | 107712 | 2 | 2048 | 20 | 310.3 | 2761 | 2.9 | 90538 |
| | 182376 | 3 | 2048 | 20 | 254.9 | 4664 | 2.9 | 126499 |
| | 437688 | 3 | 2048 | 20 | 236.1 | 3379 | 3.2 | 85122 |
| | 519876 | 2 | 2048 | 20 | 233.7 | 1957 | 3.6 | 48815 |
| | 624276 | 4 | 2048 | 20 | 229.9 | 3511 | 3.4 | 86233 |
| | 693000 | 2 | 2048 | 20 | 234.0 | 1752 | 3.7 | 43764 |
| | 777312 | 2 | 2048 | 20 | 237.0 | 1809 | 3.7 | 45739 |
| | 1058400 | 2 | 2048 | 20 | 230.0 | 830 | 3.7 | 20388 |
| | 1218960 | 2 | 2048 | 20 | 213.2 | 1465 | 3.2 | 33468 |
| | 1296000 | 3 | 2048 | 20 | 221.0 | 2109 | 3.5 | 49869 |
| ***Lyz*** | 300 | 3 | 2048 | 20 | 1436.6 | 437 | 3.1 | 38771 |
| | 1800 | 2 | 2048 | 20 | 364.7 | 632 | 3.4 | 18882 |
| | 3600 | 2 | 2048 | 20 | 881.6 | 750 | 3.1 | 35831 |
| | 7200 | 2 | 2048 | 20 | 1273.2 | 610 | 3.0 | 41984 |
| | 14400 | 2 | 2048 | 20 | 1103.4 | 519 | 3.1 | 35986 |
| | 28800 | 2 | 2048 | 20 | 1014.9 | 713 | 3.0 | 45954 |
| | 86400 | 2 | 2048 | 20 | 612.0 | 1500 | 3.5 | 79606 |
| | 172800 | 2 | 2048 | 20 | 333.6 | 2007 | 2.8 | 65634 |
| | 346600 | 2 | 2048 | 20 | 242.5 | 2592 | 2.9 | 64844 |
| | 432000 | 2 | 2048 | 20 | 211.3 | 4020 | 2.7 | 88812 |
| | 604800 | 1 | 2048 | 20 | 172.9 | 4270 | 2.8 | 78168 |
| | 1123200 | 2 | 2048 | 20 | 92.5 | 3419 | 2.6 | 35582 |
| | 600 | 3 | 2048 | 20 | 959.5 | 1402 | 3.6 | 69144 |
| | 5400 | 2 | 2048 | 20 | 591.9 | 1019 | 3.4 | 42686 |
| | 18000 | 2 | 2048 | 20 | 720.0 | 1125 | 3.0 | 55008 |
| | 91800 | 1 | 2048 | 20 | 448.8 | 541 | 3.9 | 20366 |
| | 171000 | 2 | 2048 | 20 | 495.9 | 1934 | 2.8 | 79284 |
| | 264600 | 3 | 2048 | 20 | 439.9 | 3423 | 3.4 | 133898 |
| | 346500 | 3 | 2048 | 20 | 351.2 | 4049 | 3.1 | 133400 |
| | 432900 | 3 | 2048 | 20 | 315.0 | 4798 | 2.8 | 145233 |
| | 518400 | 2 | 2048 | 20 | 313.9 | 3263 | 3.5 | 100009 |
| | 610200 | 1 | 2048 | 20 | 275.7 | 1452 | 3.5 | 39803 |
| | 688500 | 1 | 2048 | 20 | 223.1 | 2100 | 3.7 | 47373 |
| | 783000 | 1 | 2048 | 20 | 207.7 | 2082 | 3.3 | 44558 |
| | 873000 | 1 | 2048 | 20 | 181.7 | 3059 | 2.0 | 57672 |
| | 1048500 | 1 | 2048 | 20 | 164.3 | 2865 | 3.2 | 49244 |
| $\beta_2m$* | 540 | 16 | 1024 | 10 | 1002.0 | 468 | 5.9 | 36896 |
| | 3300 | 8 | 1024 | 10 | 746.8 | 515 | 5.4 | 29583 |
| | 8280 | 6 | 1024 | 10 | 616.2 | 650 | 5.5 | 29184 |
| | 16920 | 4 | 1024 | 10 | 506.6 | 603 | 5.7 | 22650 |

| | | | | | | | |
|---|---|---|---|---|---|---|---|
| 39240 | 4 | 1024 | 10 | 380.2 | 859 | 4.6 | 25747 |
| 84240 | 4 | 1024 | 10 | 301.7 | 1037 | 5.9 | 26177 |
| 108000 | 4 | 1024 | 10 | 266.0 | 1298 | 4.8 | 28612 |

* Reanalysis of data from Xue and Radford 2013 [35]

† Indicating scan size in µm x µm and image size in pixels x pixels as image aspect ratio was 1 throughout.

◊ Total number of fibril particles quantified for constructing the fibril length distributions.

° Total number of pixel height values in the fibril height distributions for fibril width evaluations.
